# ccTCM: a quantitative component and compound platform for promoting the research of traditional Chinese medicine

**DOI:** 10.1101/2023.08.04.551143

**Authors:** Dongqing Yang, Zhu Zhu, Qi Yao, Cuihua Chen, Feiyan Chen, Ling Gu, Yucui Jiang, Lin Chen, Jingyuan Zhang, Juan Wu, Xingsu Gao, Junqin Wang, Guochun Li, Yunan Zhao

## Abstract

Traditional Chinese medicine (TCM) databases play a vital role in bridging the gap between TCM and modern medicine, as well as in promoting the popularity of TCM. Elucidating the bioactive ingredients of Chinese medicinal materials is key to TCM modernization and new drug discovery. However, one drawback of current TCM databases is the lack of quantitative data on the constituents of Chinese medicinal materials. Herein, we present ccTCM, a web-based platform designed to provide a component and compound-content-based resource on TCM and analysis services for medical experts. In terms of design features, ccTCM combines resource distribution, similarity analysis, and molecular-mechanism analysis to accelerate the discovery of bioactive ingredients in TCM. ccTCM contains 273 Chinese medicinal materials commonly used in clinical settings, covering 29 functional classifications. By searching and comparing, we finally adopted 2043 studies, from which we collected the compounds contained in each TCM with content greater than 0.001%, and a total of 1 449 were extracted. Subsequently, we collected 40767 compound-target pairs by integrating multiple databases. Taken together, ccTCM is a versatile platform for that can be used by TCM scientists to perform scientific and clinical TCM studies based on quantified ingredients of Chinese medicinal materials. ccTCM is freely accessible at http://www.cctcm.org.cn.

## INTRODUCTION

As an alternative to modern medicine, traditional Chinese medicine (TCM) has been used to treat and prevent various diseases over thousands of years, playing an important role in improving the health of East Asian people [1]. In recent decades, great efforts have been exerted to study all aspects of TCM, such as clinical evaluation [2], chemical profiling [3], and bioactivities [4]. With the rapid increase in available TCM data, many web-based databases specializing in TCM have emerged, which in turn facilitate the scientific and clinical study of TCM.

First, the TCM Information Database [5], published in 2006, has been introduced earlier as a web resource to provide free-of-charge and comprehensive information about TCM, including herbs, prescriptions, herbal ingredients, structure and functional properties of compounds, as well as their therapeutic effects and clinical indications and applications. The database represents early efforts toward enhancing the ability to evaluate TCM herbs’ beneficial and risk effects.

Second, are TCM Database@Taiwan [6] and SymMap [7], which emphasize phenotypic drug discovery (PDD) based on a large amount of information on natural products and their clinical applications. TCM Database@Taiwan, published in 2011, contains more than 20,000 natural compounds from 453 Chinese Materia Medica, including herbs, animal products, and minerals. The 2D and 3D formats of each compound in the database are available for virtual filtering or molecular simulation. SymMap, an integrative database of TCM enhanced by symptom mapping, was published in 2019. SymMap presents the newly curated symptom-herb knowledge and connects symptoms and phenotypes to herbs and diseases, thereby providing both phenotypic changes and lead compounds for PDD screening efforts.

Finally, for TCMID [8], TCMSP [9], HERB [10], and LTM-TCM [11], these four TCM databases focus on understanding of the action mechanisms underlying TCM through the concept and theory of network pharmacology. TCMID, a TCM integrative database for herb molecular mechanism analysis, was published in 2012. Based on predicted targets of compounds, the database displays herb-disease networks and compound-target networks, integrating TCM with modern science at the phenotypic and molecular levels. TCMSP, a database of system pharmacology for drug discovery from herbal medicines, was published in 2014. The database improves the network-pharmacology analysis of TCM with the help of absorption, distribution, metabolism, and excretion related properties of compounds. Unlike TCMID and TCMSP, HERB, published in 2020, used gene targets guided by high-throughput transcriptomic screening experiments to identify herb-disease networks and compound-target networks. LTM-TCM, published in 2022, is currently the most comprehensive TCM database. Using the LTM-TCM platform, the network-pharmacology analysis of TCM is enhanced by large amounts of data integration and high-quality normalization.

Nowadays, these TCM databases play a crucial role in bridging the gap between TCM and modern medicine as well as in promoting the modernization and popularization of TCM [12]. However, some problems have emerged in the result reliability of PDD screening and network-pharmacology analysis. An obvious disadvantage of these TCM databases is that all of them do not provide quantitative data on ingredients in Chinese medicinal materials. Some ingredients of Chinese medicinal materials with very low content, even if they have better bioactivities, are not responsible for the therapeutic effects of TCM medicinal materials. If such ingredients are not discarded based on content data, the ranking of lead compounds and the construction of herb-compound-target-disease networks are bound to be seriously affected. Thus, adding quantitative data on ingredients into TCM databases contributes to upgrading PDD screening and network-pharmacology analysis.

With the rapid development of quantitative analysis techniques including chemical and instrumental analysis methods [13], many studies have aimed to determine the contents of different ingredients of Chinese medicinal materials and link them to the biodiversity and quality evaluation of Chinese medicinal materials [14]. In particular, a few TCM quality-control studies have focused on detecting the content differences of multiple compounds in certain Chinese medicinal materials derived from different botanical origins [15], producing areas [16], cultivation years [17], harvesting seasons [18], and processing methods [19] through high-performance liquid chromatography (HPLC) coupled with different up-to-date detectors. The rapidly increasing number of quantified ingredients in Chinese medicinal materials provides us an opportunity to develop a component and compound-content-based database integrating comprehensive information on TCM (**ccTCM**).

Hence, in the present study, we obtained the quantified ingredients data of TCM based on comprehensive literature searches with focus on botanical origins, producing areas, harvesting seasons, and processing methods. The scope of our collection covered 29 categories of TCM, 273 Chinese medicinal materials (Supplementary Table S1), and 1499 compounds in total. The content-determination information of each TCM was obtained through manual literature retrieval, totaling 2043 articles, with an average of 7.48 literature supporting each TCM. For the convenience of use, the metadata of each TCM was collected from the Chinese Pharmacopoeia (2020 edition), and the metadata of each compound was collected from PubChem [20]. We also provided 40 767 pieces of target information for compounds. In brief, the data-based connections between TCM and modern medicine described in ccTCM provided reliable support for understanding the molecular mechanisms underlying TCM clinical therapy. Moreover, ccTCM provided similarity analysis of Chinese medicinal materials and resource-distribution analysis of components and compounds, thereby enabling the progress of the TCM industry and scientific research.

## METHODS

### Data sources of Chinese medicinal materials and compounds

The metadata of Chinese medicinal materials in ccTCM (including name, species, Latin name, medicinal part, basic characteristics, dosage, toxicity, main efficacy, identification, source information, trait, storage conditions, etc.) originated from the Chinese Pharmacopoeia (2020 edition) and were automatically translated into English through Google translation API.

The metadata of compounds was retrieved, using the compound name and their synonyms, from PubChem by using PubChemPy (v1.0.4) package via PubChem’s PUG REST web service. The metadata of each compound includes molecular formula, molecular weight, complexity, classification, properties, synonyms, IUPAC, InChi, InChiKey, Canonical Smiles, Isomeric Smiles, exact mass, etc. The structures of compounds were searched from PubChem, ChEMBL [21], and ZINC [22]. For those compounds not found in these databases, their structures were drawn by using InDraw 5.2 software (https://www.integle.com/static/indraw).

### Manual collection of quantified TCM ingredients

The schematic of document retrieval, quantitative data collection, and ingredient rating is shown in Figure 1. The chemical profiling of Chinese medicinal materials was searched from the Chinese Academic Journal Network Publishing Database (CAJD) (https://www.cnki.net/) by using the combinations of the keywords (“name”, “progress”, and “chemistry”) in the title. Original research on ingredient content analysis was also searched from CAJD by using the combinations of the keywords (“name”, “content”, and “determination”) in the title. The 5777 pieces of literature 273 Chinese medicinal materials were adopted, as the botanical origin of Chinese herbs and the content unit of quantified ingredients was clearly clarified in the context. The inclusion criteria of ingredient content data into ccTCM can be referring to supplementary document 1. In brief, quantitative data were preferentially extracted from the articles focusing on the quality assessment of Chinese medicinal materials derived from different botanical origins, producing areas, cultivation years, harvesting seasons, or processing methods through the quantitative analysis of multi-components by a single marker or HPLC-based simultaneous detection. Abnormal quantitative data that deviated so far from the rest of the data were not adopted.

**Figure 1.**
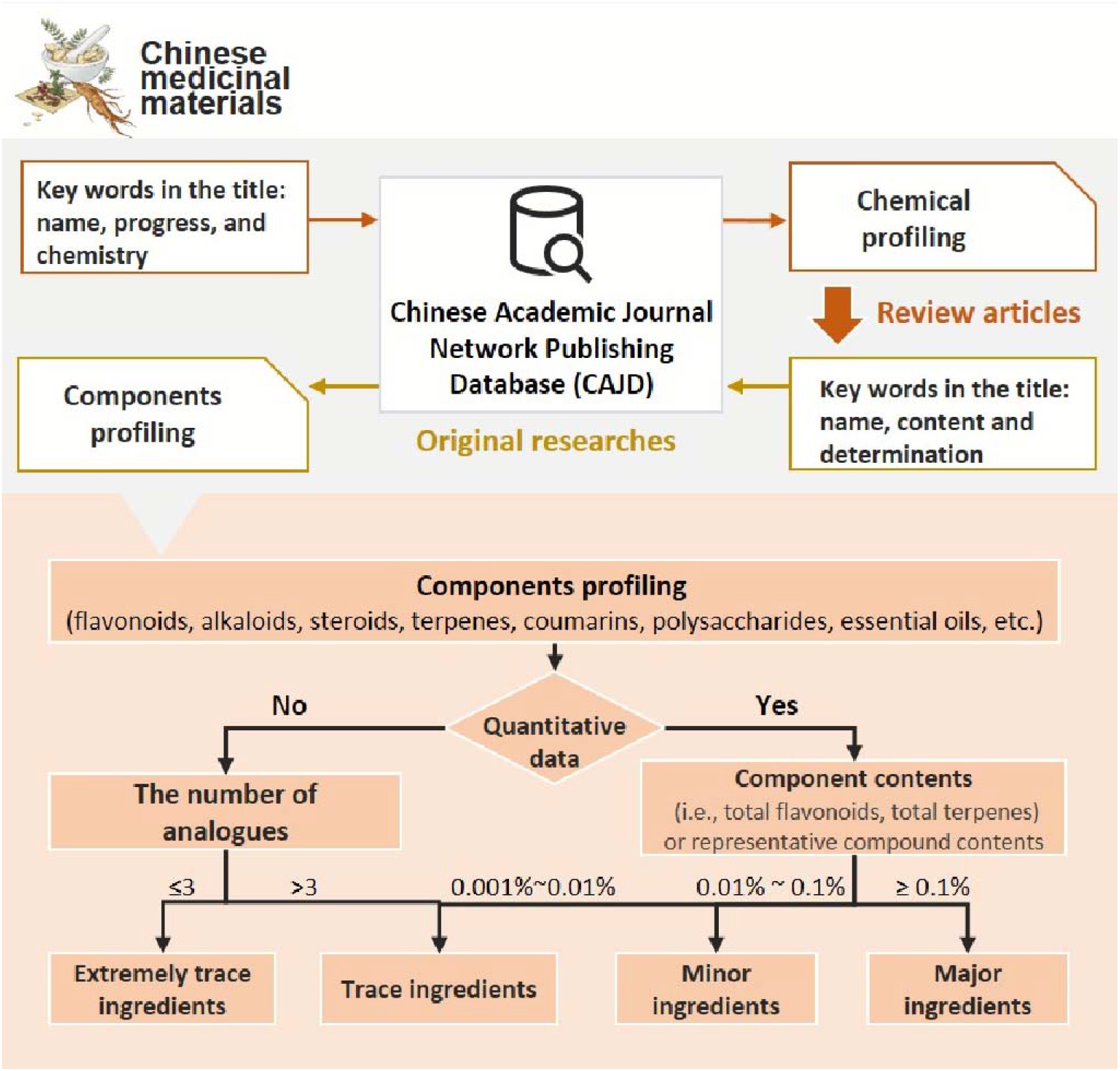
Schematic of document retrieval, quantitative data collection, and ingredient rating. Related papers were collected through Chinese Academic Journal Network Publishing Database (CAJD). These articles were selected as candidates, in the context of which the botanical origin of Chinese herbs and the content unit of quantified ingredients is clearly clarified. The inclusion criteria of ingredient content data into ccTCM can be referring to supplementary document 2. The basis for the setting of major ingredients was as follows: if the patient was given 10 g of medicinal materials per day, the value of 0.1% (g/g) indicated that the patient can take 10 mg of ingredients. In fact, most of drugs are orally used at a dosage of not less than 10–20 mg per day. Particularly, alkaloids were identified as major ingredients if the content of total alkaloids or a representative compound exceeded 0.01%. When the content of polysaccharides, aliphatic organic acids or fatty oils in medicinal materials exceeded 10%, they can be considered as major ingredients.

We divided ingredients into 26 categories based on the structural characteristics of natural products, which were named major category in ccTCM, including aliphatic organic acids, alkaloids, benzyls, caffeoylquinic acids, chromones, coumarins, diarylheptanoids, essential oils, fatty oils, flavonoids, inorganic compounds, lignans, nucleosides, phenanthrenes, phenols, phenylethanols, phenylpropanoids, polyacetylenes, polypeptides, polysaccharides, quinones, steroids, stilbenes, tannins, terpenes, and others. Each major category contained some minor categories and subcategories, with a total of 115 minor categories and 132 subcategories (Supplementary Table S2). These ingredients were regarded as major ingredients when component contents (i.e., total flavonoids and total terpenes) or representative compound contents were equal to or greater than 0.1% (g/g). Minor and trace ingredients were defined as component contents or representative compound contents of 0.01%–0.1% (g/g) and 0.001%–0.01% (g/g), respectively. If no quantitative data existed, such ingredients that possessed more than three analogs were regarded as trace ingredients. As regards the weight factor of ingredients in TCM, we ranked major ingredients, minor ingredients, and trace ingredients as 1, 0.3, and 0.1.

### Compound–target relationships

We collected compound-target relationships primarily by integrating multiple reliable databases, such as Human Metabolome Database (HMDB, v5.0) [23], DrugBank v5.1.9 [24], Comparative Toxicogenomics Database (CTD, 2022-04) [25], Natural Product Activity and Species Source (NPASS v2022) [26], and Collective Molecular Activities of Useful Plants (CMAUP, v1.0) [27]. To avoid omission of information, we used compound names and their synonyms for matching. The literature links of compound primarily**–**target were provided when the PubMed IDs were available.

### Implementation of ccTCM

The ccTCM database was developed on the PostgreSQL database (v14.0) and Django server framework (v3.2). Its web interfaces were built using the Vue3 framework, and ECharts was used for front-end visualization. The entire database was designed to enable the access of its entries by TCM and compounds by using multiple browse and search facilities. When applicable, the compound entries were cross-linked to the PubChem, CTD, and ZINC databases. The relevant pieces of literature on compound**–**target relation was provided by PubMed identifiers and cross-linked to PubMed. ccTCM is freely accessible at http://www.cctcm.org.cn without a need for user registration. The website is compatible with most major browsers. Enrichment analysis was conducted using the R package “clusterProfiler” (v4.2.2) (Yu et al. 2012), and networkx (https://networkx.org/, v2.6.3) was used for network-module analysis and net-properties calculations including diameter, clustering coefficient, closeness centrality, and betweenness centrality.

## RESULTS

### Database statistics

ccTCM currently contains 273 Chinese medicinal materials containing 1,449 unique compounds targeting 9,880 proteins. We collected a total of 1,248 records of TCM component or representative compound contents with 1,073 supporting literature, a total of 2,757 TCM-compound content pairs with 1,126 supporting literature, and 40,767 compound**–**target pairs (Table 1).

**Table 1.** Overview of data from peering databases.

| Items | Data type | ccTCM | TCMSP | TCMID 2.0 | ETCM | HERB | HIT 2.0 |
| --- | --- | --- | --- | --- | --- | --- | --- |
| Published year |  |  | 2014 | 2018 | 2019 | 2020 | 2022 |
| Chinese | The total number | 273 | 499 | 8159 | 402 | 7263 | 1237 |
| medicinal | Origin images | Yes | No | n.a. | No | No | No |
| materials | Identification images | Yes | No | n.a. | Yes | No | No |
| Compounds | The total number | 1449 | 29 384 | 43 413 | 7284 | 49 258 | 1284 |
|  | Classification | Yes | No | n.a. | No | No | No |
|  | Druglikeness | Yes | No | n.a. | Yes | No | No |
|  | ADMET properties | Yes | Yes | n.a. | Yes | No | No |
| Quantification | Quantified ingredients | 1248 | 0 | 0 | 0 | 0 | 0 |
| information | TCM-compound content pairs | 2757 | 0 | 0 | 0 | 0 | 0 |
| Targets | The total number | 9880 | 3311 | 82 | 2266 | 12 933 | 2208 |
|  | Source by literatures supported | Yes | No | n.a. | No | Yes | Yes |
|  | Compound-target pairs | 40767 | 84260 | n.a. | n.a. | 4815 | 10 031 |
| Literature | The total number | 7027 | 1288 | n.a. | n.a. | 1966 | 7100 |
| Online<br>analysis | Resource distribution and similarity analysis | Yes | No | No | No | No | No |
|  | Molecular-mechanism analysis (network analysis and enrichment analysis (KEGG and GO)) | Yes | No | No | Yes | Yes | No |

n.a.: not available.

On the ccTCM main page, users can view the sunburst plot containing all Chinese medicinal materials and click the tick next to the TCM name to open the detail page. These Chinese medicinal materials were classified into 7 categories according to TCM function, and each category was further divided into 29 subcategories (Supplementary Table S1).

### Browsing and searching Chinese medicinal materials, compounds, and literature

Users can view all Chinese medicinal materials, compounds, and literature through the resource browser. The TCM category filter can help users screen the list of Chinese medicinal materials. Similarly, compound browsing can also be filtered by a major category filter. Users can specify a range of years to view the list of available literature.

The resource browser also provides different angles for users to view the data contained in ccTCM. The component profile lists the weight factors of components included in each TCM by using the numbers 1, 0.3, and 0.1. In the component content page, users can view the content data of components or representative compounds in each TCM, and each record provided the corresponding literature. The compound-content page lists the quantified compounds in each TCM, whose average contents in Chinese medicinal materials are generally more than 0.01% (g/g). Each compound was given a structural classification including major category, minor category, and subcategory.

The search page is convenient for users to search for wanted Chinese medicinal materials, compounds, and targets included in ccTCM. The search keywords can be the names of Chinese medicinal materials or compounds in English or Chinese.

On each TCM page, users can visit the metadata, origin picture, identification pictures, component profiling, compound contents, and corresponding targets on which the compounds act (Figure 2). On each compound page, users can visit the molecular formula, molecular weight, complexity, classification, properties, cross-references and corresponding targets (Figure 3).

**Figure 2.**
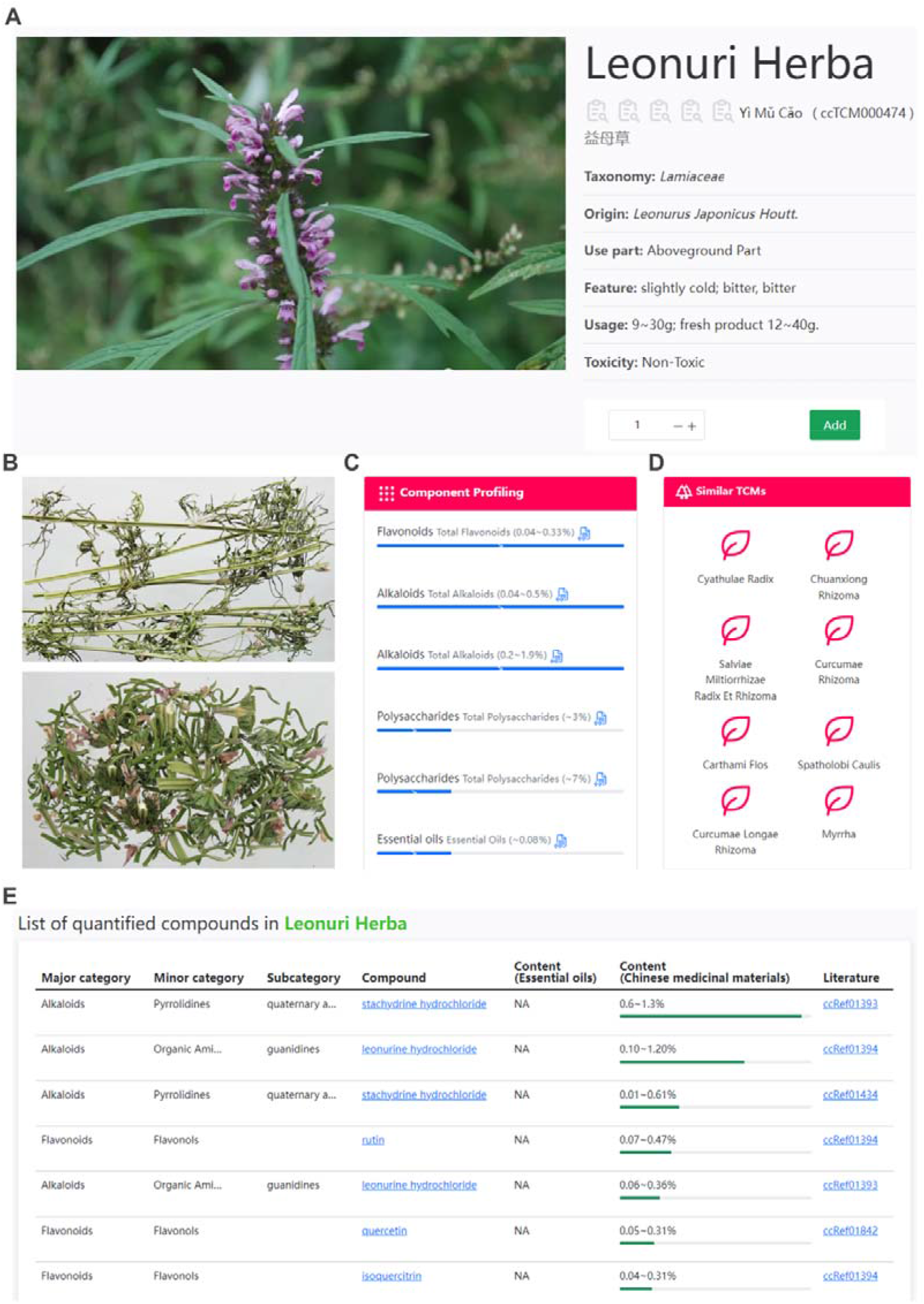
The metadata of Leonurus japonicas is taken as an example (A): taxonomy, origin, medicinal part, feature, usage, and toxicity. (B) The identification image of the morphology of L. japonicas. (C) Component profiling of L. japonicas. (D) Lists of similar TCMs belonging to the same functional category. (E) List of all the quantified compounds in L. japonicas with a content ratio greater than 0.001%.

**Figure 3.**
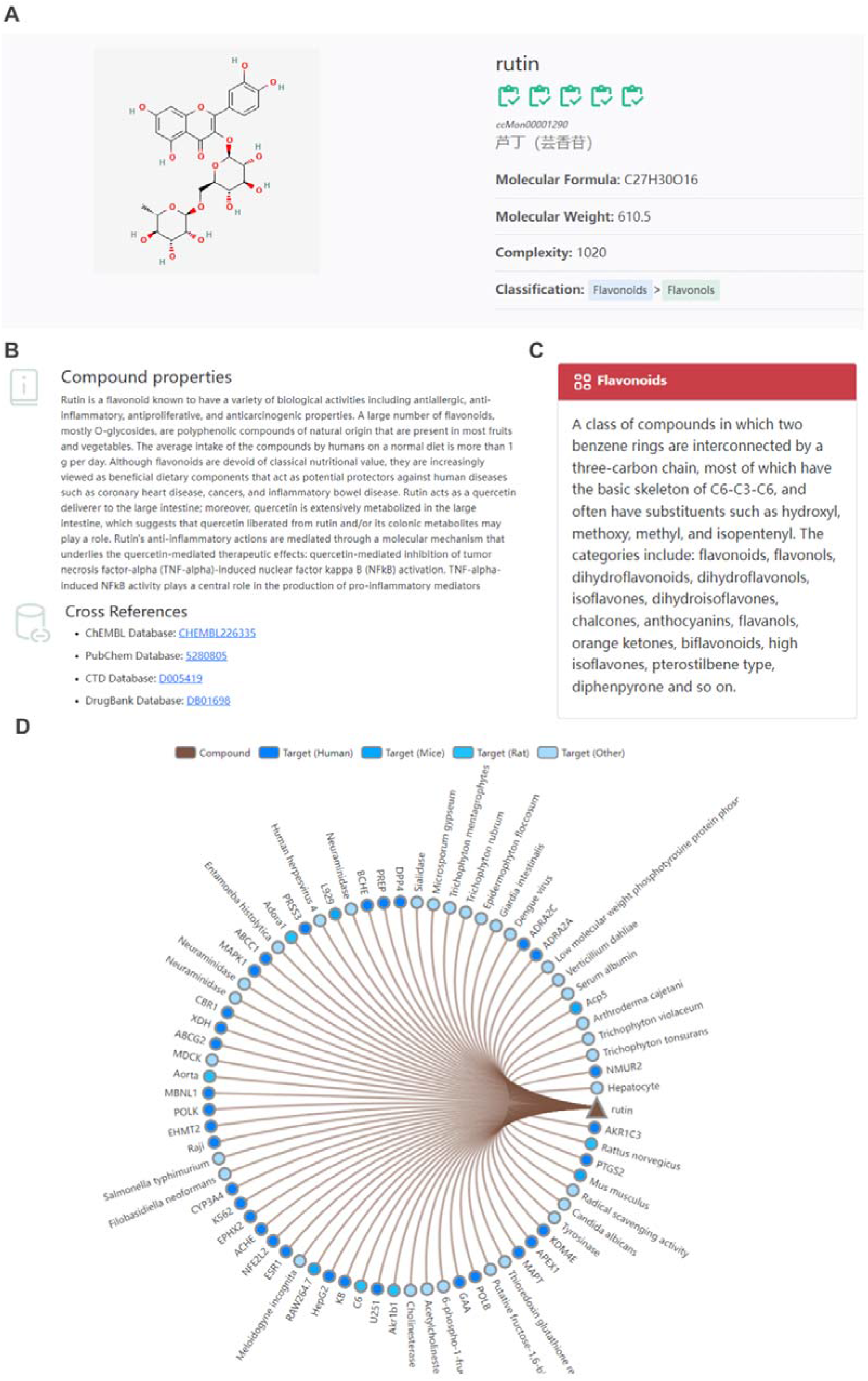
Information of a specific compound. (A) The metadata of rutin is taken as an example: molecular formula, molecular weight, complexity, and classification. (B) Description of compound properties and available cross-references. (C) Functional description of the main category to which the compound rutin belongs. (D) Network of relation between rutin and targets in humans, mice, rats, and other organisms.

### Pot function, take Gegen Qinlian Tang as an example

Gegen Qinlian Tang (GQT) is mostly used in diarrhea and diabetes clinically. This prescription contains 15 g of Puerariae Lobatae Radix, 9 g of Scutellariae Radix, 9 g of Coptidis Rhizoma, and 6 g of Glycyrrhizae Radix Et Rhizoma. ccTCM provides a Pot function like a shopping cart for users to customize the prescription on their own. On the Pot page, the prescription can be named, and the TCM quantity can also be modified or even deleted (Figure 4A). At the bottom of the page, users can view all the compounds and their quantitative information contained in the current prescription (Figure 4B). To demonstrate the reliability of the quantitative information provided by ccTCM, we compared it with the measurement data in [28], and the comparison results are shown in Table 2. Spearman’s rank correlation analysis showed that the quantitative data provided by ccTCM and the data measured by Li et al had high consistency (*r*=0.943, *P* value=0.005). The details of the formulation using Pot function and subsequent molecular mechanism analysis can be referring to supplementary document 2.

**Figure 4.**
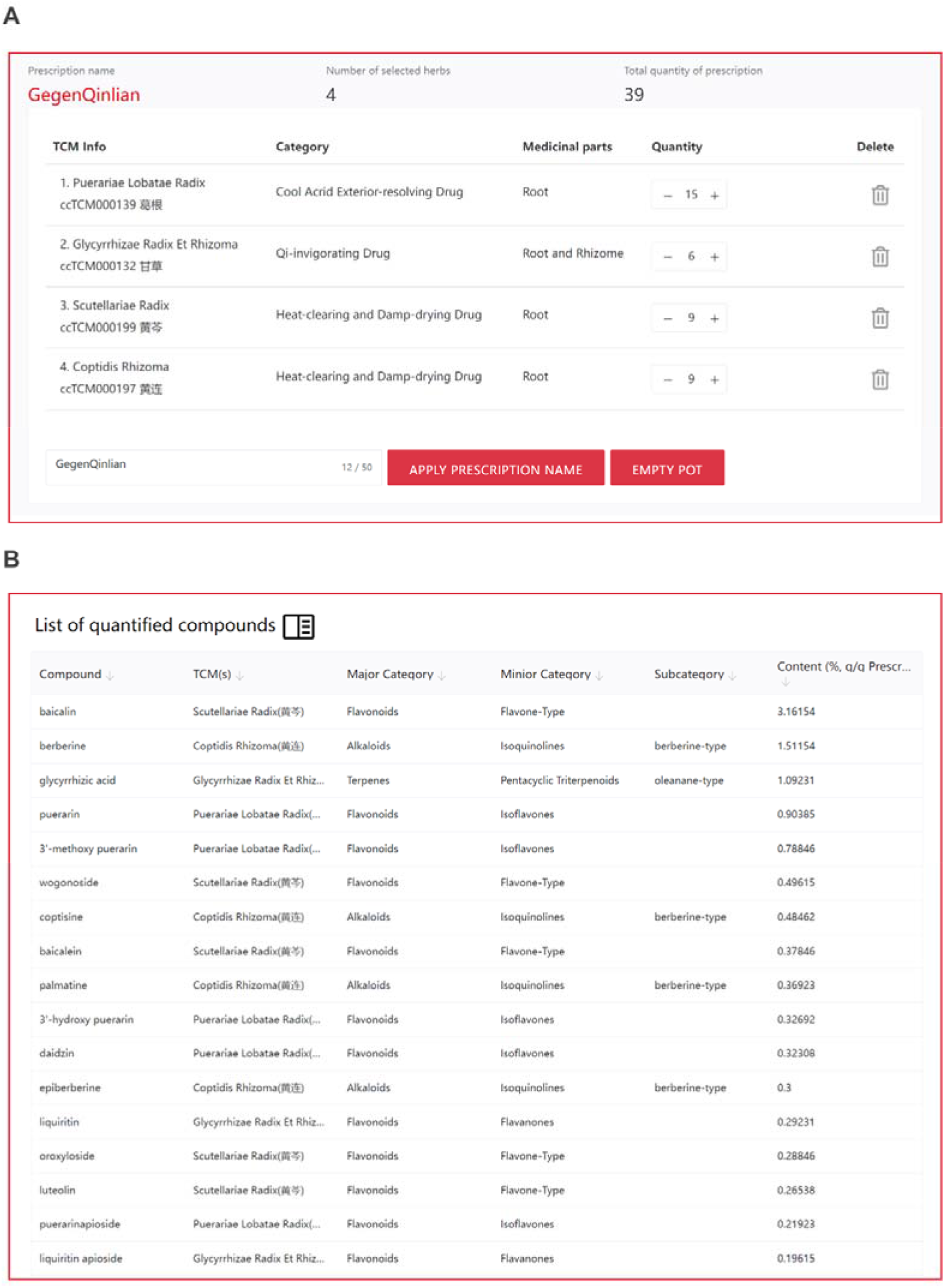
Pot function. (A) In the pot function, users can customize the quantity of various Chinese medicinal materials. (B) All TCM compounds and their quantitative information contained in the current prescription.

**Table 2.** Comparison of quantitative information of compounds in GQT.

| Compound | Source | Content<br>in GQT extract<br>(%)<br>[28] | Median content<br>(ccTCM)<br>(%, g/g<br>Prescription) | Content<br>(ccTCM)<br>(%, g/g<br>Prescription) |
| --- | --- | --- | --- | --- |
| Puerarin | *Pueraria lobata (Willd) Ohwi.* | 2.87 | 0.90385 | 0.115~1.692 |
| Daidzein | *Pueraria lobata (Willd) Ohwi.* | 0.77 | 0.01538 | 0.0038~0.0269 |
| Liquiritin | *Glycyrrhiza uralensis Fisch* | 1.42 | 0.29231 | 0.05~0.54 |
| Baicalin | *Scutellaria baicalensis*<br>*Georgi.* | 28.84 | 3.16154 | 2.17~4.15 |
| Baicalein | *Scutellaria baicalensis*<br>*Georgi.* | 9.72 | 0.37846 | 0.04~0.72 |
| Berberine | *Coptis chinensis Franch.* | 18.93 | 1.51154 | 1.2~1.8 |

### Resource distribution, similarity analysis, and molecular-mechanism analysis

The resource distribution of ingredients in Chinese medicinal materials can be viewed by selecting or specifying compounds or components. ccTCM uses cascade mode to facilitate users to select the component object (Figure 5A). For example, the user wants to find the distribution data of oxindole-type alkaloids in Chinese medicinal materials. First, alkaloids in the drop-down box of the major category are selected, and then indoles in the drop-down box of minor category are selected. Finally, the oxindole-type alkaloid in the drop-down box of subcategory is selected. Users can directly type the name of the compound in the dialog box to view the distribution in Chinese medicinal materials (Figure 5B).

**Figure 5.**
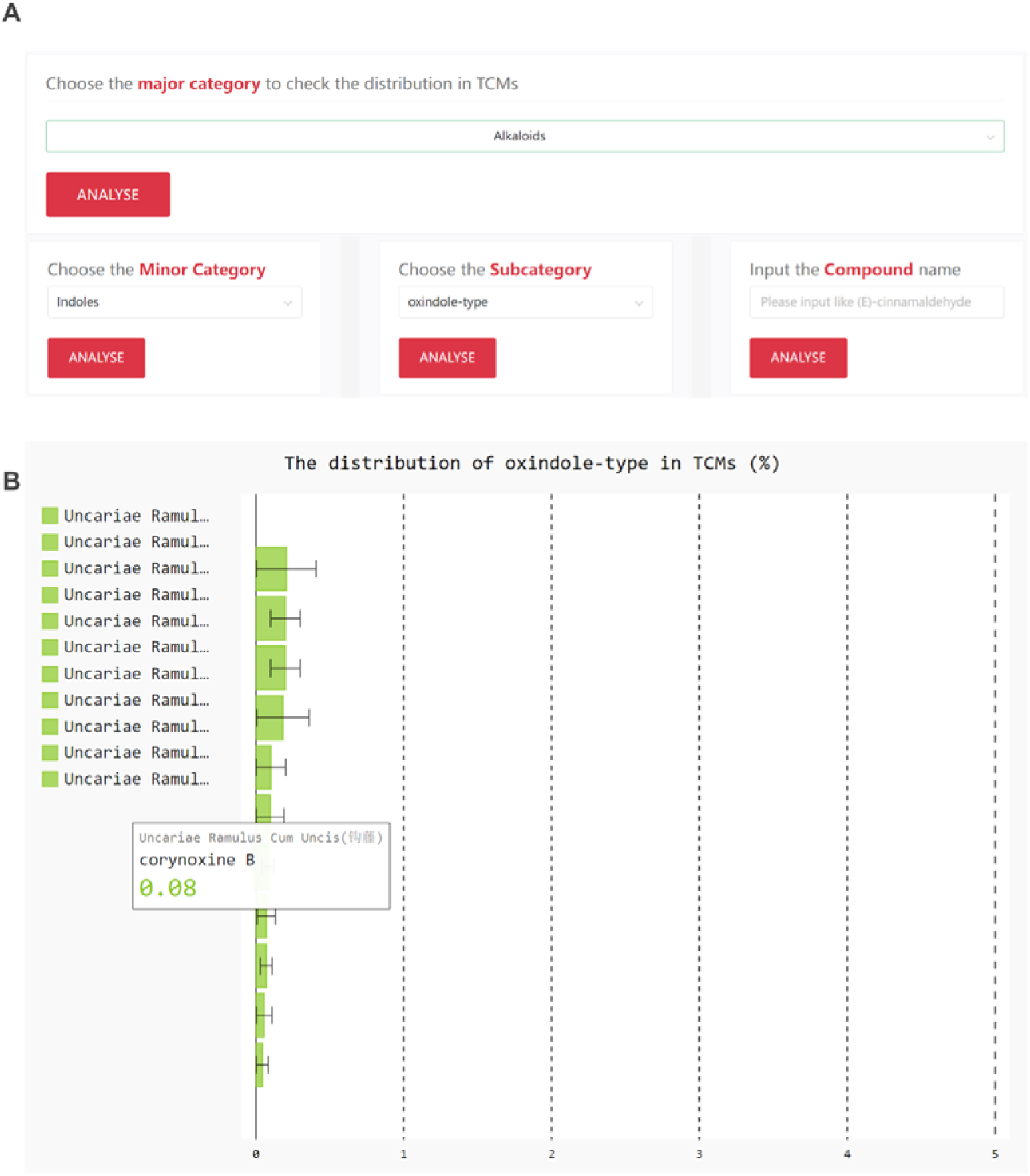
Presentation of resource-distribution analysis. (A) The drop-down menu provides users with resource analysis at different aspects (major category, minor category, and subcategory) in a cascading manner. (B) Bar plot of distribution of Bisepoxylignans in ccTCM.

The TCM similarity analysis service provides a comparison of Chinese medicinal materials from three aspects: major category, minor category, and compound. By using the Pot function, users add TCM to the Pot and specify the quantity. The analysis method uses Spearman’s rank correlation coefficient, and the analysis results are displayed in the form of heat map (Figure 6A).

**Figure 6.**
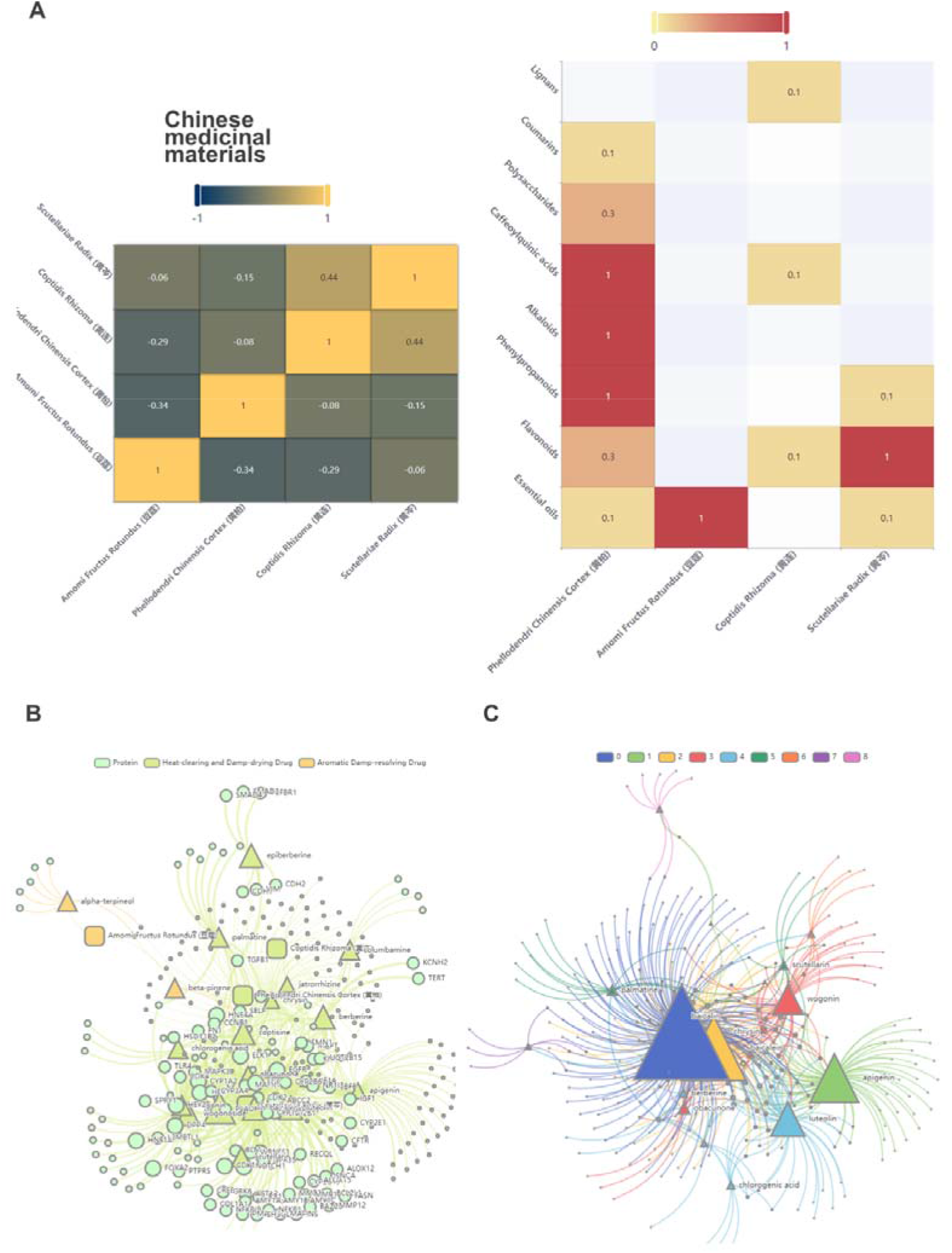
Presentation of enrichment-analysis results. (A) Similarity analysis uses a similarity matrix to reflect the Spearman correlation among different Chinese medicinal materials. A heatmap showing the contents of compounds in each TCM. (B) Weighted compound-target network, with the nodes resized according to their degrees. (C) Module-identified network is analyzed according to the network module-identification algorithm, and different colors represent different possible modules.

Molecular-mechanism analysis refers to network analysis and enrichment analysis (KEGG signaling-pathway enrichment analysis and gene ontology (GO) functional-module enrichment analysis) according to the quantified compounds in Chinese medicinal materials and the targets they act on. The currently accepted species are *Homo sapiens*, *Mus musculus*, and *Rattus norvegicus*. ccTCM provides three types of networks (Compound Target Network, Weighted Compound Target Network, and Module Identified Network).

As for the Compound Target Network, the box represents TCM, the triangle represents compound, and the circle represents gene. Different colors represent different classifications of Chinese medicinal materials or compounds. The TCM node size corresponds with its quantity in the prescription, and the compound node size corresponds with its quantity in the TCM multiplied by the TCM quantity in the prescription. As regards the Weighted Compound Target Network, the nodes were resized according to their degrees, which was an update of the previous network. A higher content of compounds in the prescription corresponded with a larger size of the compound nodes and corresponding gene nodes connected to them (Figure 6B). The Module Identified Network is analyzed according to the network-module identification algorithm [29], and different colors represent different possible modules (Figure 6C). All analysis results are available for user download.

### Special subject of COVID-19

For the treatment of COVID-19, China has accumulated a considerable clinical experience in the aspect of TCM therapy and has proposed many effective prescriptions. The ccTCM platform provided the three prescriptions (Qingfei Paidu Decoction, Huashi Baidu Prescription, and Xuanfei Baidu Prescription) suggested by the State Administration of Traditional Chinese Medicine (http://www.satcm.gov.cn/xinxifabu/meitibaodao/2020-04-17/14712.html, Supplementary Table S3). Users can easily view the contents of the three prescriptions from the home page and carry out molecular-mechanism analysis. We also marked effective traditional Chinese materials for COVID-19 treatment on TCM pages.

## DISCUSSION AND CONCLUSION

Accurate quantitative information plays a crucial role in expediting the discovery of effective ingredients in TCM and its formulations, thereby promoting the development of novel drugs. While there has been a rapid accumulation of quantitative data from laboratory and clinical studies on TCM herbs and ingredients in recent decades, there has been a lack of a well-structured organizational system to catalog this information. Furthermore, the latest TCM-related references published in the past decade have remained uncurated. Consequently, this study aimed to address these gaps by meticulously gathering all available literature pertaining to the determination of the quantity of TCM ingredients and curating high-confidence target information from recently published TCM references. Leveraging the Pot function and online analysis, we have successfully constructed ccTCM, the sole database encompassing quantitative information for all TCM compounds currently available.

The novelty of the ccTCM database includes the following: (i) ccTCM is the first available database containing quantitative component and compound data in Chinese medicinal materials; (ii) ccTCM integrates the Pot function for the user-defined analysis of molecular mechanism of TCM, visualized-distribution profiles of components and compounds in Chinese medicinal materials, and similarity analysis of different Chinese medicinal materials from three aspects (major category, minor category, and compounds). (iii) ccTCM is the first available database providing structural classification of natural compounds. The current version of ccTCM contains a total of 273 Chinese medicinal materials and covers almost all functional classifications of TCM. Nevertheless, some Chinese medicinal materials have not been collected yet. We plan to add more Chinese medicinal materials into the ccTCM database in the future. We will also try to integrate TCM theory into ccTCM and provide a more comprehensive and useful TCM database.

## ACKNOWLEDGMENTS

The authors thank Weichao Xu for authorizing ccTCM to use his TCM images. The study was financially supported by National Natural Science Foundation of China (No.82204647, 82003970), and supporting project of National Natural Youth Foundation of Nanjing University of Chinese Medicine (XPT82204647), and supporting project of Jiangsu Province "The 14th Five-year Plan" Key Discipline-Public Health and Preventive Medicine (035091005007).

## AUTHOR CONTRIBUTIONS

Yunan Zhao and Dongqing Yang designed the research. Zhu Zhu, Qi Yao, Cuihua Chen, Feiyan Chen, Ling Gu, Yucui Jiang, and Lin Chen collected and corrected the data. Dongqing Yang designed the ccTCM database and developed the website. Jingyuan Zhang, Juan Wu, and Xingsu Gao tested the website performance and correction. Junqin Wang and Guochun Li performed the statistical analyses. Yunan Zhao and Dongqing Yang wrote the manuscript.

## DATA AVAILABILITY STATEMENT

All data provided by ccTCM is accessible for free at http://www.cctcm.org.cn/.

## ORCID

Yunan Zhao http://orcid.org/0000-0003-4944-8765

## Supplementary materials

Table S1 List of Chinese medicinal materials information contained in ccTCM

Table S2 Classification information list of TCM compounds

Table S3 Prescription list for COVID-19

