## supplementary document 1 for "ccTCM: a quantitative component and compound platform for promoting the research of traditional Chinese medicine"

Inclusion criteria of ingredient quantitative information into ccTCM

Example 1: take hyperoside in Apocyni Veneti Folium as an example

| No. | Content (%)^*^ | Literature |
| --- | --- | --- |
| 1 | 0.21~0.29 | Zhou Chunling, Sun Lingling, Bi Kaishun. RP-HPLC method for the determination of hyperoside and apocytin in Apocynum venetum leaves. Journal of Pharmaceutical Analysis, 2009, 29 (6): 1001-1003.  (周春玲，孙苓苓，毕开顺. RP-HPLC法测定罗布麻叶中金丝桃苷和罗布麻甲素的含量. 药物分析杂志，2009, 29(6): 1001~1003) |
| 2 | 0.05~0.32 | Liu Xunhong, Zhang Yuechan, Li Junsong, et al. Simultaneous determination of four flavonoids in Apocynum venetum leaves by HPCE-DAD. Chinese Journal of Pharmacy, 2010, 45 (6): 464-467.  (刘训红，张月婵，李俊松，等. HPCE-DAD 同时测定罗布麻叶中 4 种黄酮的含量. 中国药学杂志，2010, 45(6): 464~467) |
| 3 | 0.26~0.32 | Zhang Qunlin, Wu Liang, Yan Anding, et al. Simultaneous determination of six flavonoids in Apocynum venetum leaves and their extracts by HPLC. Chinese Journal of Traditional Chinese Medicine, 2011, 36 (5): 589-593.  (张群林，吴亮，言安定，等. HPLC 同时测定罗布麻叶药材及其提取物中6种黄酮的含量. 中国中药杂志，2011, 36(5): 589~593) |
| 4 | 0.27~0.36 | Xu Shuo, Xu Wenfeng, Kuang Yongmei, et al. The content of six flavonoid components in Apocynum venetum leaves was determined using a one test multiple evaluation method. Journal of Pharmaceutical Analysis, 2019, 39 (7): 1217-1228  (徐硕，徐文峰，邝咏梅，等. 一测多评法测定罗布麻叶中 6 个黄酮类成分的含量. 药物分析杂志，2019, 39(7): 1217~1228) |
| 5 | 0.06~0.15 | Chen Shuai, Wang Liang, Qi Rongrong, et al. Simultaneous determination of 7 components in Apocynum venetum leaves using UPLC method. Traditional Chinese patent medicines and simple preparations, 2020, 42 (12): 3211~3215.  (陈帅，王亮，齐绒绒，等. UPLC 法同时测定罗布麻叶中7 种成分. 中成药，2020, 42(12): 3211~3215) |
| 6 | 0.40~0.50 | Wu Shu-chen, He Minyou, Li Guowei, et al .UPLC characteristic spectra and flavonoid content determination of Apocynum venetum leaves from different regions. Chinese Modern Traditional Chinese Medicine, 2021, 23 (4): 619-626.  (吴淑珍，何民友，李国卫，等. 不同产地罗布麻叶 UPLC 特征图谱及黄酮类成分含量测定. 中国现代中药，2021, 23(4): 619~626) |

* The Chinese Pharmacopoeia 2020 edition stipulates that the content of hyperoside in Apocyni Veneti Folium shall not be less than 0.3%. Using the limited content in the pharmacopoeia as a reference, the content data measured in literature 2, 4, and 6 were included in the database (0.05-0.50%). Literature 1, 3, and 5 were not included in the database as their content data fall within the above range.

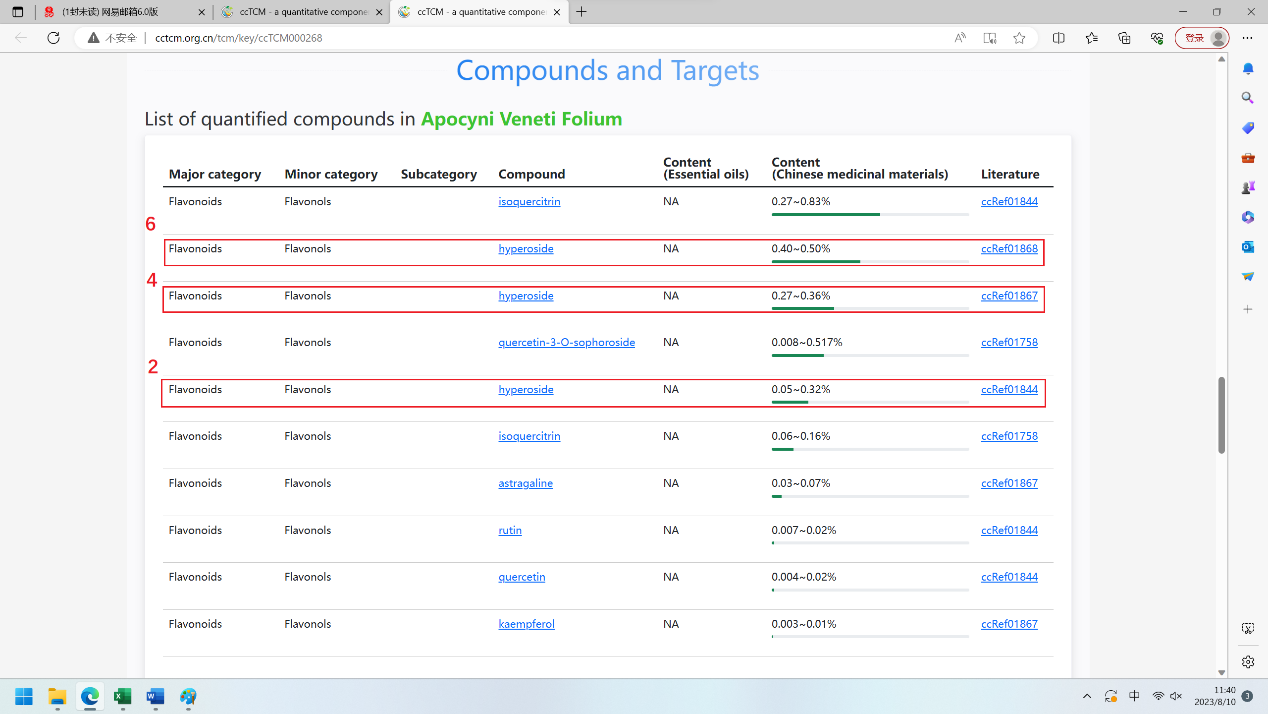

Example 2: Take formononetin in Spatholobi Caulis as an example

| No. | Content (%)^*^ | Literature |
| --- | --- | --- |
| 1 | 0.002~0.012 | Zheng Yan, Wang Bin, Wang Jingli, et al. HPLC determination of Flavonoid in Spatholobus spatholobi from different habitats. Chinese Journal of Traditional Chinese Medicine, 2008, 33 (15): 1920-1922  (郑岩，王邠，王京丽，等. 高效液相测定不同产地鸡血藤药材中黄酮类化合物的含量. 中国中药杂志，2008, 33(15): 1920~1922) |
| 2 | 0.003~0.021 | Li Ying, Chen Xiaohui, Zhang Tianhong, et al. RP-HPLC method for the determination of anthocyanin in Caulis spatholobi. Journal of Shenyang Pharmaceutical University, 2009, 26 (12): 975~977  (李莹，陈晓辉，张天虹，等. RP-HPLC法测定鸡血藤中芒柄花素的含量. 沈阳药科大学学报，2009, 26(12): 975~977) |
| 3 | 0.004~0.037 | Liang Yongshu, An Ran, Liu Junmin, et al. Determination of Genistein and Stigmarin in Caulis Spatholobi from different habitats. Shizhen Guoyi Guoyao, 2013, 24 (7): 1655-1657  (梁永枢，安冉，刘军民，等. 不同产地鸡血藤药材中染料木素及芒柄花素的含量测定. 时珍国医国药，2013, 24(7): 1655~1657) |
| 4 | 0.001~0.019 | Chen Hongying, Yan Ping, Zhang Min, et al. High performance liquid chromatography fingerprint and analysis of anthocyanin content in Caulis spatholobi from different origins. Journal of Guangzhou University of Chinese Medicine, 2015, 32 (5): 923~928  (陈红英，严萍，张敏，等. 不同产地鸡血藤药材高效液相指纹图谱及芒柄花素含量分析. 广州中医药大学学报，2015, 32(5): 923~928) |
| 5 | 0.019~0.023 | Chen Qianping, Gu Xiaoyu, Long Hairong, et al. Determination of Mangostensin and Total Flavonoids in Caulis spatholobi from Different Producing Areas. Contemporary Chemical Industry, 2016, 45 (7): 1549-1552  (陈乾平，谷筱玉，龙海荣，等. 不同产地鸡血藤药材中芒柄花素及总黄酮的含量测定. 当代化工，2016, 45(7): 1549~1552) |

* The 2020 edition of the Chinese Pharmacopoeia does not have a limit on the content of formononetin in Spatholobi Caulis. Taking multiple content values as a reference, the content data measured in literature 3 and 4 were included in the database (0.001~0.037%). Literature 1, 2, and 5 were not included in the database as their content data fall within the above range.

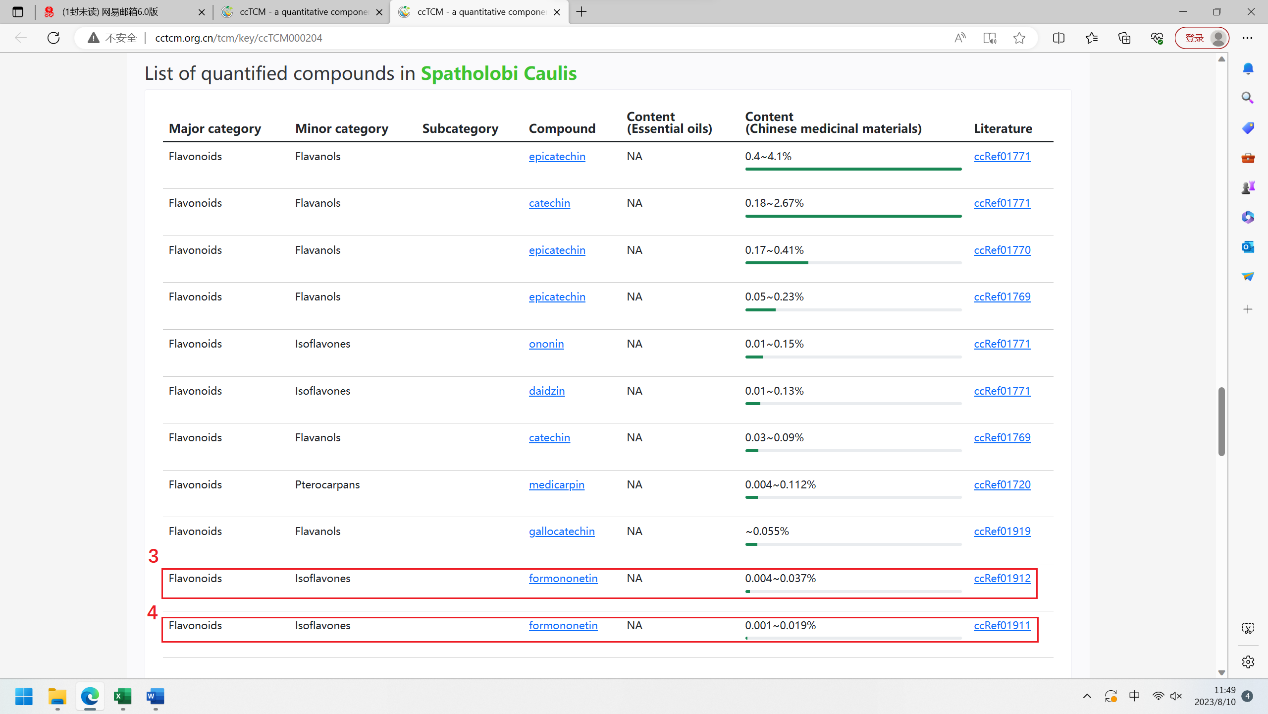

Example 3: take amygdalin in Persicae Semin as an example

| No. | Content (%)^*^ | Literature |
| --- | --- | --- |
| 1 | 0.01~0.04 | Wang Youlan, Li Hongbing, Hua Yuqin. HPLC method for determining the content of amygdalin in peach kernel. Chinese Pharmacist, 2002, 5 (9): 550, 556  (王友兰，李红兵，华玉琴. HPLC法测定桃仁中苦杏仁苷的含量. 中国药师，2002, 5(9): 550, 556) |
| 2 | 2.2~3.5 | Pan Huichao, Qin Wenhong. HPLC method for determining the content of amygdalin in bitter almonds and peach kernels. World's Latest Medical Information Digest, 2004, 3 (5): 1321-1322  (潘会朝，秦文红. HPLC法测定苦杏仁和桃仁中苦杏仁甙的含量. 世界最新医学信息文摘，2004, 3(5): 1321~1322) |
| 3 | 2.4~3.3 | Xiao Xiong, Wei Huizhen, Zhang Dan, et al. HPLC determination of D-amygdalin in bitter almonds, peach kernels, and Yuli kernels. Journal of Jiangxi University of Traditional Chinese Medicine, 2019, 31 (3): 76~79  (肖雄，魏惠珍，张丹，等. HPLC 测定苦杏仁、桃仁、郁李仁中D-苦杏仁苷的含量. 江西中医药大学学报，2019, 31(3): 76~79) |
| 4 | 1.7~3.9 | Zhang Xuelan, Zhang Zhipeng, Deng Lihong, et al. Study on the Differences in Content and Characteristic Maps of Peach Kernel and Its Processed Products. Chinese herbal medicine, 2019, 42 (12): 2803~2808  (张雪兰，张志鹏，邓李红，等. 桃仁及其炮制品含量和特征图谱差异性研究. 中药材，2019, 42(12): 2803~2808) |
| 5 | 1.5～4.7 | Zhang Congcong, Wang Changhong, Li Xingjia, et al. Based on UHPLC-Q-Orbitrap HRMS, analyze the chemical components of peach kernels and quickly determine the content of amygdalin and kurarin. Chinese Journal of Medicine and Pharmacy, 2022, 42 (4): 347-355  (张聪聪，王长虹，李兴佳，等. 基于 UHPLC-Q-Orbitrap HRMS分析桃仁化学成分及快速测定苦杏仁苷和野黑樱苷的含量. 中国医药药学杂志，2022, 42(4): 347~355) |

* The 2020 edition of the Chinese Pharmacopoeia stipulates that the content of amygdalin in Persicae Semin shall not be less than 1.5%. Using the limited content in the pharmacopoeia as a reference, the content data measured in literature 5 were included in the database (1.5-4.7%). Reference 2, 3, and 4 were not included in the database as their content data fall within the above range. The data measured in literature 1 deviated significantly and were not included in the database.

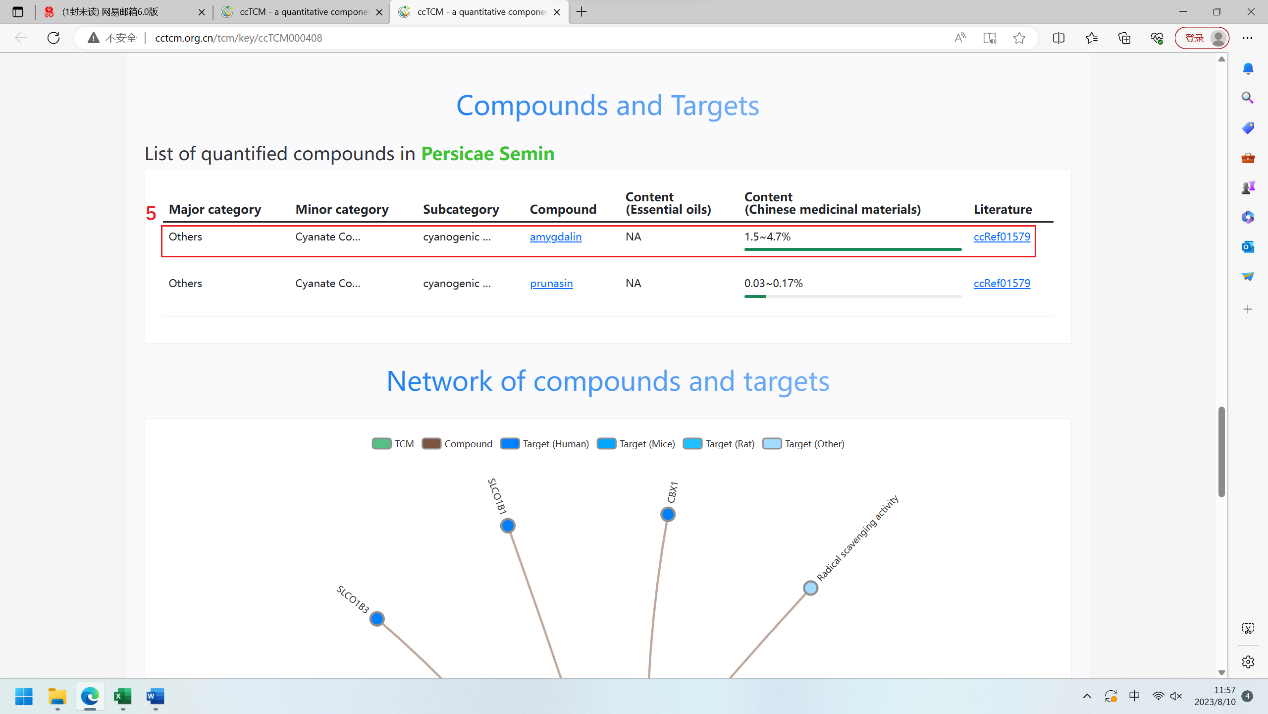

Example 4: Inclusion criteria of compound relative content data in essential oils (take ar-turmerone in essential oils of Curcumae Longae Rhizoma as an example)

| No. | Relative Content (%)^*^ | Literature |
| --- | --- | --- |
| 1 | 12.7~29.9 | Zhang Ping, Zhang Guizhi, Fan Qingyue, et al. GC-MS analysis of the characteristic chemical components of volatile oil in turmeric slices. Modern Traditional Chinese Medicine Research and Practice, 2008, 22 (3): 41-44  (张萍，张桂芝，樊晴月，等. GC-MS法分析姜黄饮片挥发油的特征性化学成分. 现代中药研究与实践，2008, 22(3): 41~44) |
| 2 | 43.53~55.99 | Yang Qing, Yan Xiaoxia, Wang Maoyuan, et al. Chemical composition and antioxidant activity of volatile oil from turmeric from different origins. Traditional Chinese patent medicines and simple preparations, 2016, 38 (5): 1188~1191  (羊青，晏小霞，王茂媛，等. 不同产地姜黄挥发油的化学成分及其抗氧化活性. 中成药，2016, 38(5): 1188~1191) |
| 3 | 33.61~49.26 | Yang Qing, Yan Xiaoxia, Wang Maoyuan, et al. GC-MS analysis of volatile oils from Hainan turmeric at different harvesting periods. Journal of Tropical Crops, 2014, 35 (9): 1866-1870  (羊青，晏小霞，王茂媛，等. 不同采收期海南姜黄挥发油的GC-MS 分析. 热带作物学报，2014, 35(9): 1866~1870) |
| 4 | 5.72 | Qiang Yueyue, Wei Hang, Fang Ling, et al. HS-SPME-GC-MS analysis of the chemical components of the volatile oil from Fujian turmeric. China Foods Limited Additives, 2020, 31 (1): 147~153  (强悦越，韦航，方灵，等. 福建姜黄挥发油化学成分的 HS-SPME-GC-MS分析. 中国食品添加剂，2020, 31(1): 147~153) |
| 5 | 8.20~25.78 | Liu Hongxing, Chen Fubei, Huang Chusheng, et al. Comparative study on two extraction methods of volatile oil from Guangxi turmeric. Guangxi Botany, 2007, 27 (5): 796-800  (刘红星，陈福北，黄初升，等. 广西姜黄挥发油两种提取方法的比较研究. 广西植物，2007, 27(5): 796~800) |
| 6 | 12.63~28.17 | Ye Shiyun, Huang Yongqi, Luo Hongmei, et al. Analysis of volatile oil from Curcuma longa in Nanbei Panjiang area of Guizhou province by gas chromatography-mass spectrometry. Agricultural science, 2012, 40 (10): 5989~5990  (叶世芸，黄勇其，骆红梅，等. 贵州南北盘江地区姜黄挥发油成分的气相色谱-质谱联用分析. 安徽农业科学，2012, 40(10): 5989~5990) |
| 7 | 0 | Tang Xiaowen, Chen Guobin. Gas chromatography-mass spectrometry analysis of the chemical components of volatile oil from turmeric. Journal of Mass Spectrometry, 2004, 25 (3): 163-165  (唐课文，陈国斌. 气相色谱-质谱法分析姜黄挥发油化学成分. 质谱学报，2004, 25(3): 163~165) |

* The 2020 edition of the Chinese Pharmacopoeia does not specify a limit on the content of ar-turmerone in essential oils of Curcumae Longae Rhizoma. In different references, the relative content of ar-turmerone in essential varies greatly (0-55.99%), which is also a common characteristic of compound content in essential oils. Therefore, for the relative content of compounds in essential oils, only the maximum attainable content data (55.99%) was included in the database.

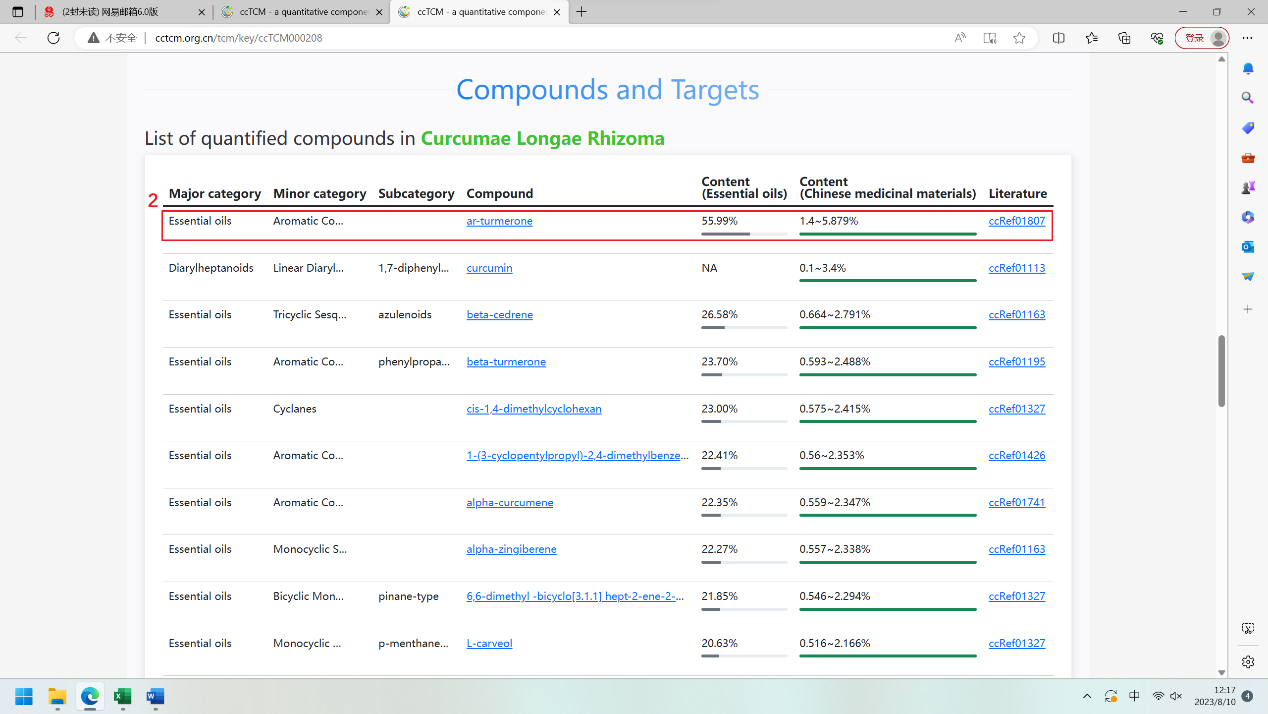
