## supplementary document 2 for "ccTCM: a quantitative component and compound platform for promoting the research of traditional Chinese medicine"

Tutorial of TCM Formulation and Molecular Mechanism Analysis Using ccTCM

**Step 1** Enter the desired traditional Chinese medicine on the search page, and click on the search result to enter the TCM page.

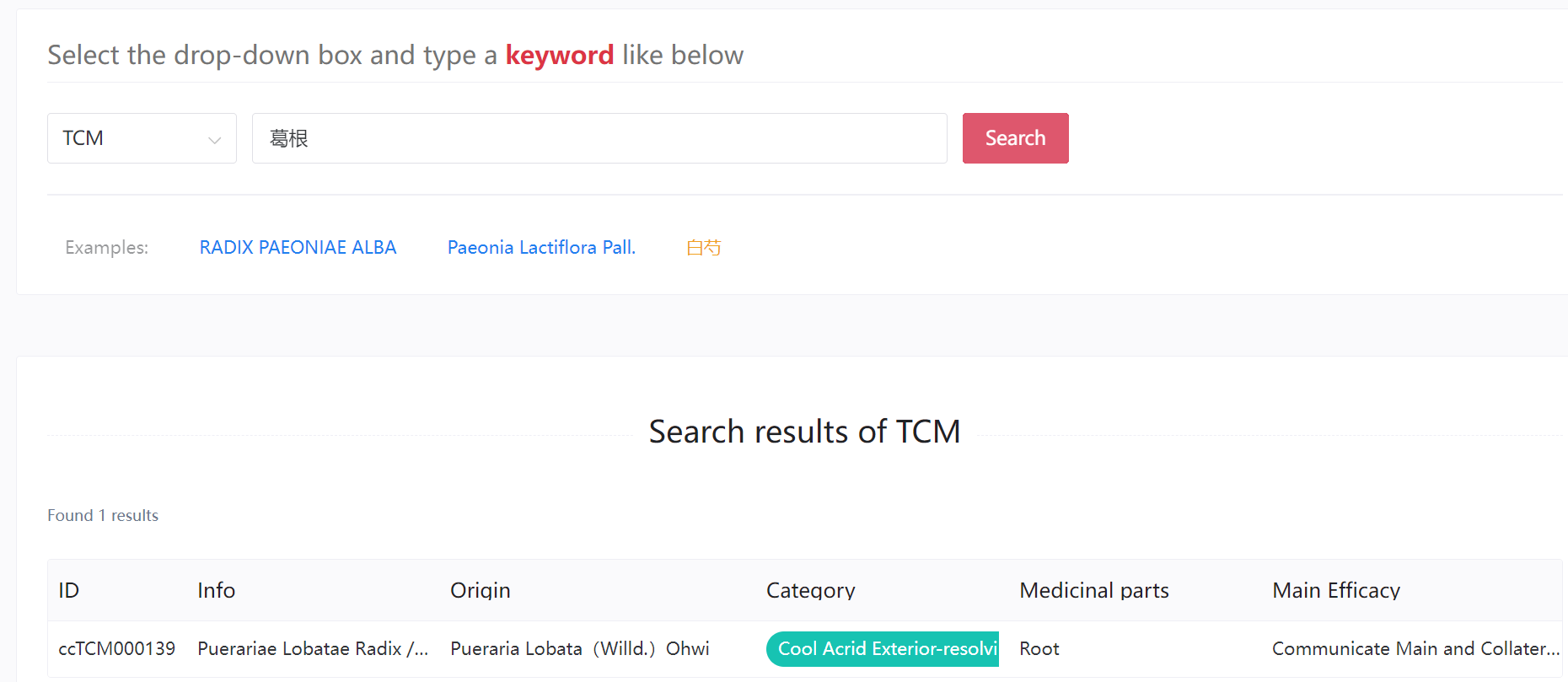

**Step 2** On the TCM page, set the quantity and click the Add button, so that the medicine will enter the Pot (upper right corner of the page).

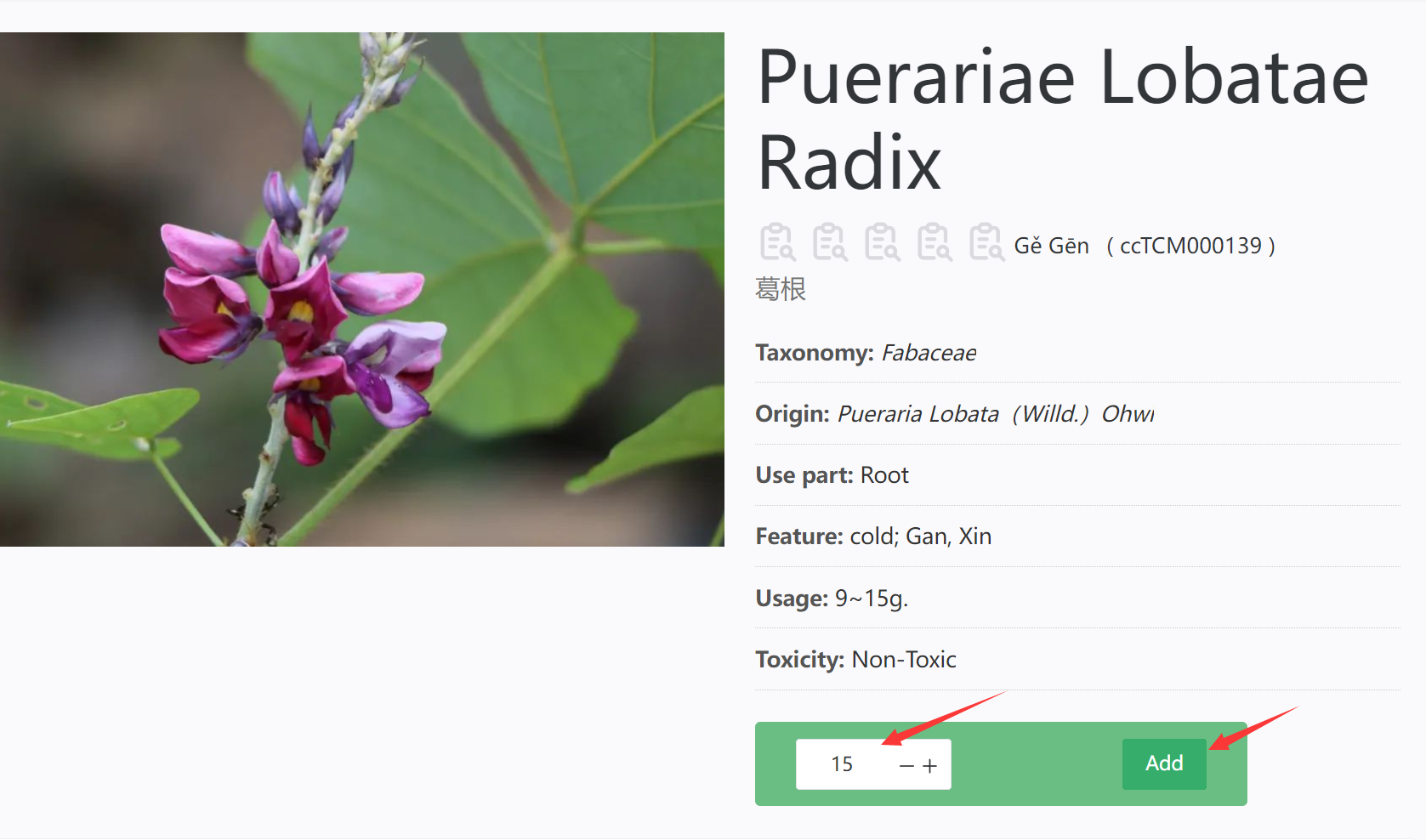

Repeat steps 1 and 2 until the required TCM is formulated according to the ratio relationship, and this information will be saved in Pot. By clicking on the Pot icon in the upper right corner of the page, in the pop-up page on the right, click Edit button to enter the Pot page.

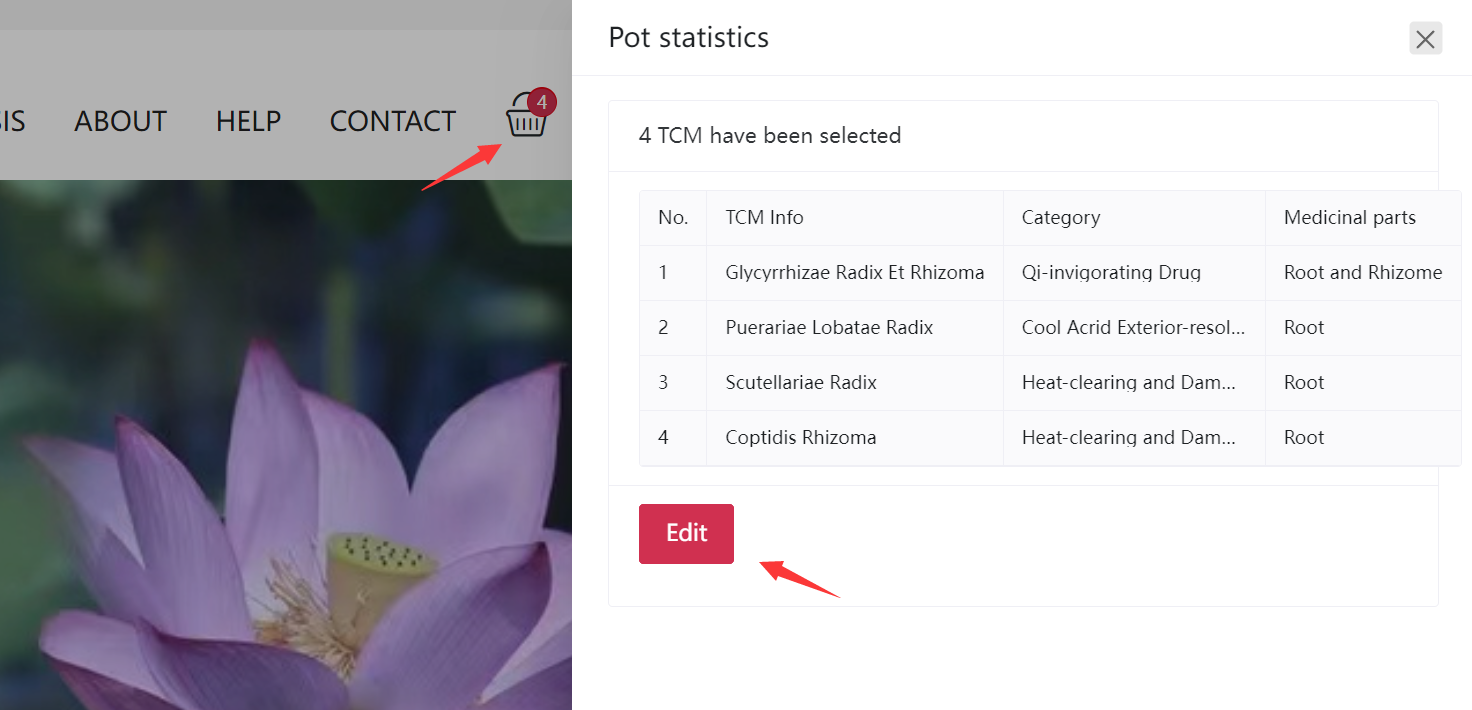

**Step 3** The users can view detailed information and make modifications on the Pot page. The prescription can be named through the text box in the bottom left corner, and the quantity of each TCM can also be modified or deleted (The data in Pot uses the cookie technology, which expire in one month. This function requires the user's browser to accept cookies.).

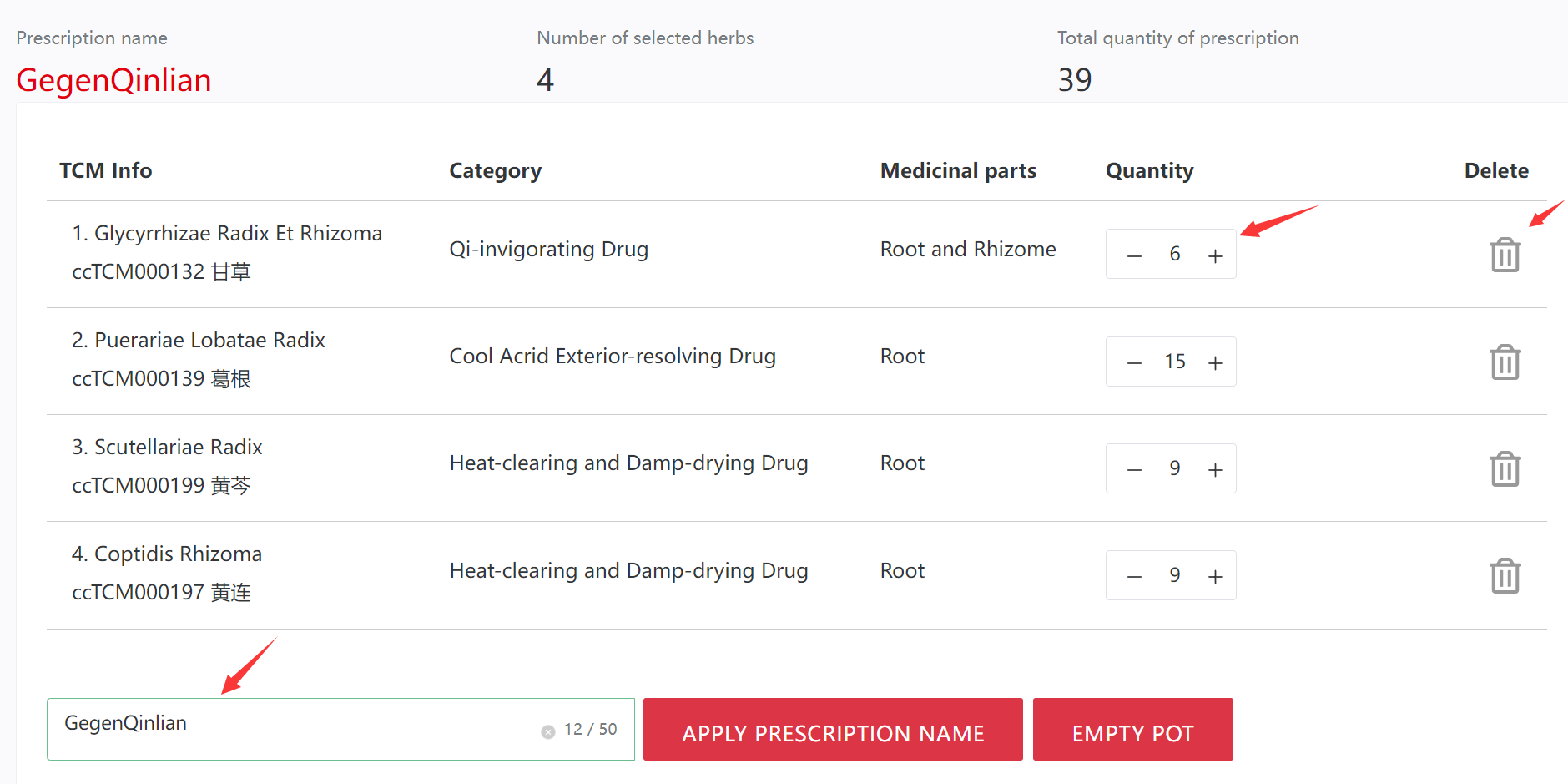

You can view the content information of all compounds in the current prescription below and find the compound with the highest content by sorting.
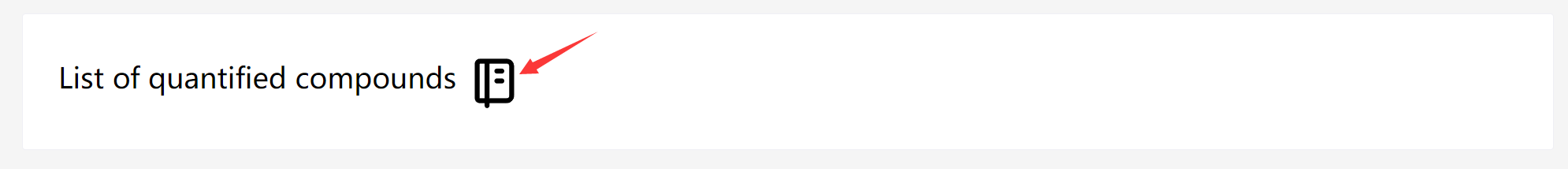

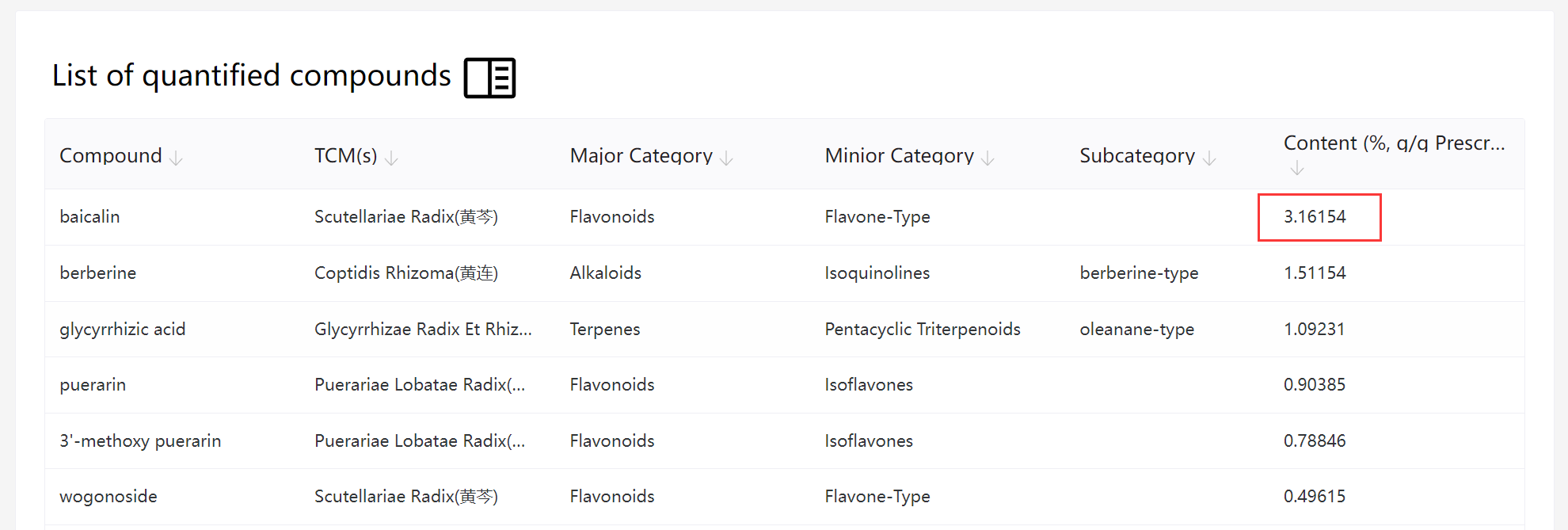

**Step 4** Enter the molecular mechanism analysis page through the menu.
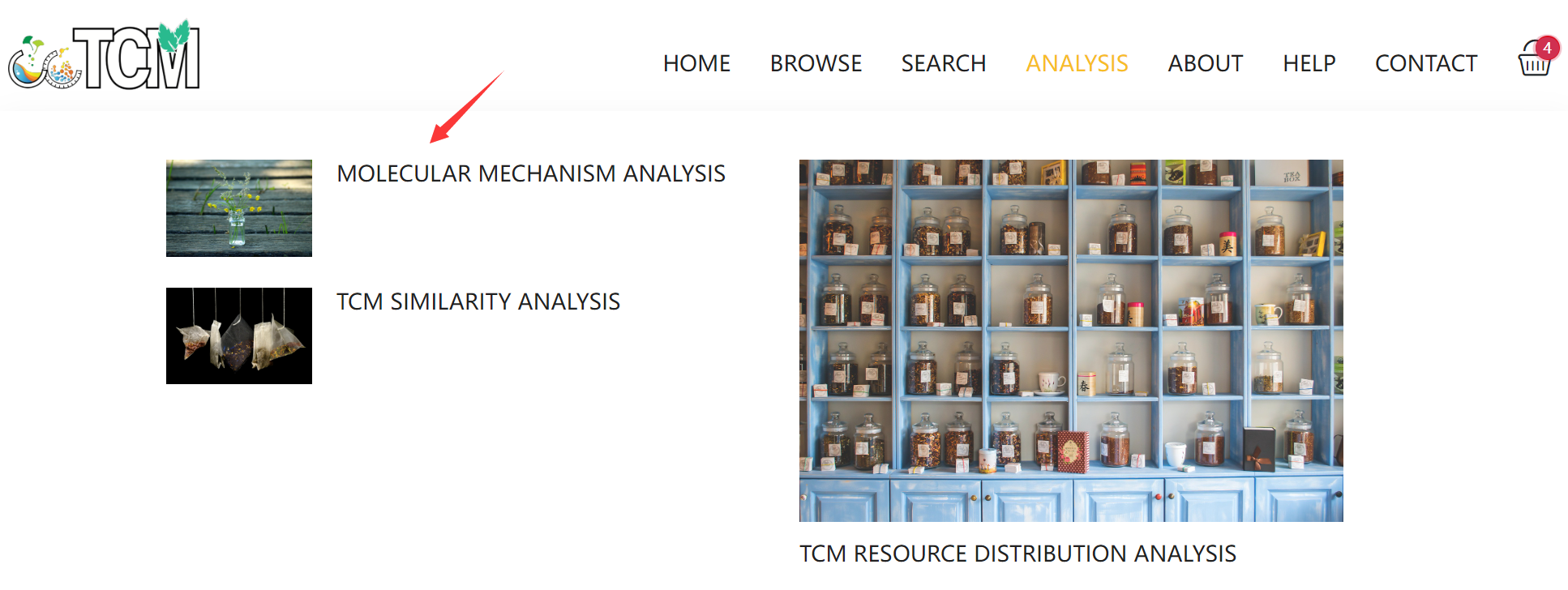

**Step 5** Select species (*Homo sapiens*, *Mus musculus*, or *Rattus norvegicus*) for molecular mechanism analysis, which mainly includes network analysis and enrichment analysis (KEGG and GO).

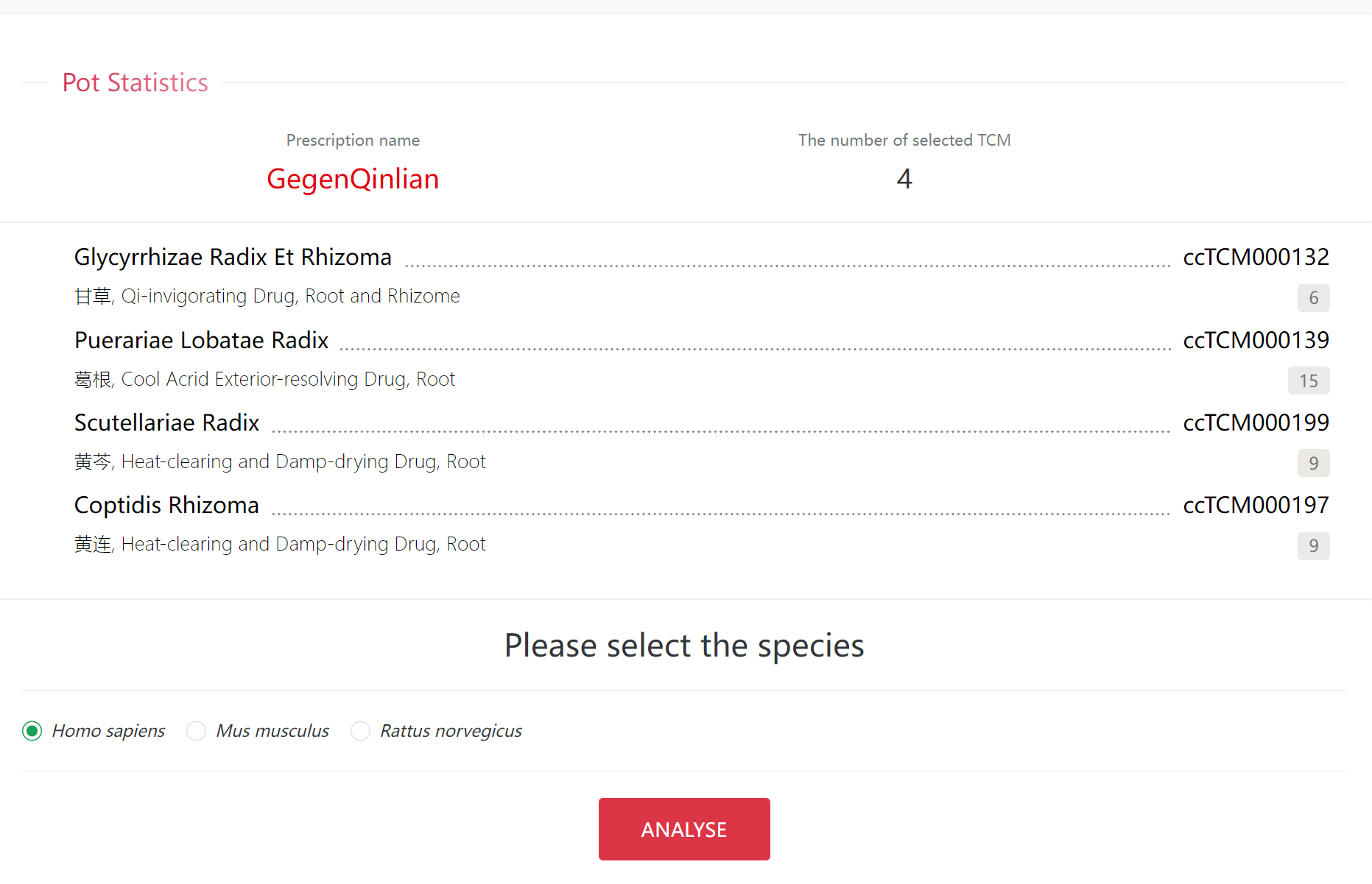

This analysis will take some time to complete, please be patient and wait.
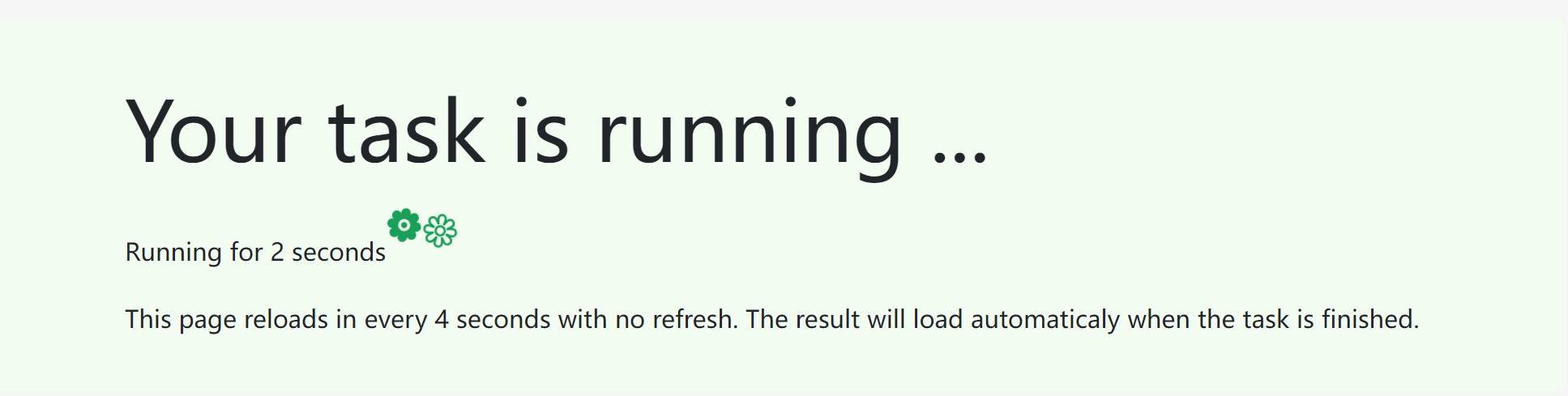

The first part of the results is network analysis. ccTCM provides three types of networks (Compound Target Network, Weighted Compound Target Network, and Module Identified Network): Compound Target Network: box represents TCM, triangle represents compound, and circle represents gene. Different colors represent different classifications of TCMs or compounds. The TCM node size corresponds with its proportion in the prescription, and the compound node size corresponds with its proportion in the TCM multiplied by the TCM proportion in the prescription.

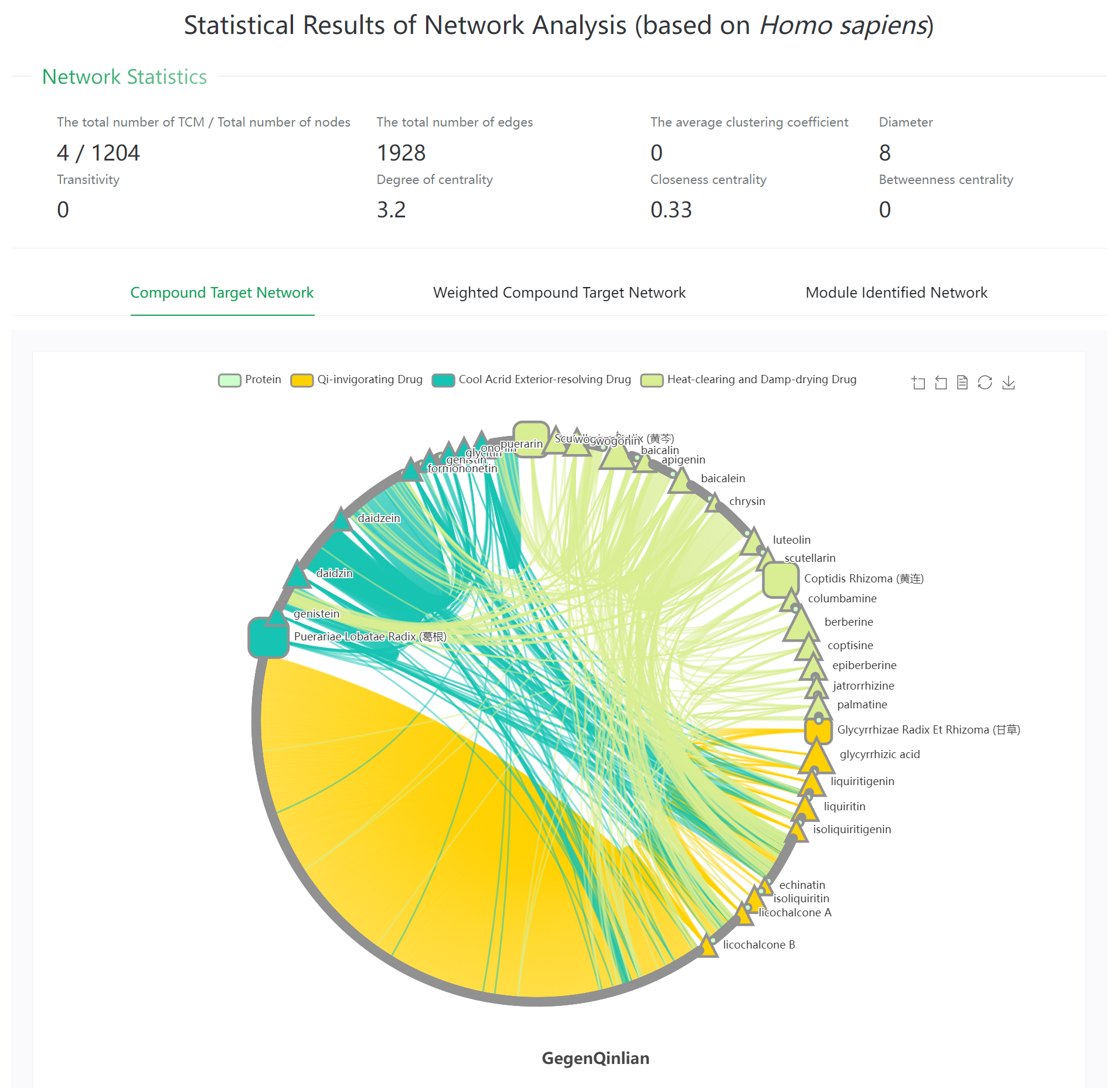

The above analysis may not meet all the needs, so users are encouraged to conduct customized analysis by downloading nodes files and links files by clicking the blue button.

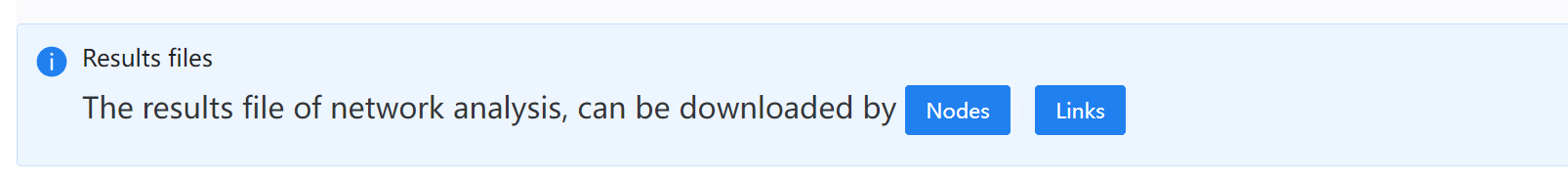

**Step 6** ccTCM also provides common functional enrichment analysis, including KEGG Pathway analysis and GO enrichment analysis (implemented by clusterProfiler). Only the top 20 records with the lowest *p* value are displayed on the page, and all the results can be viewed by clicking Download button.

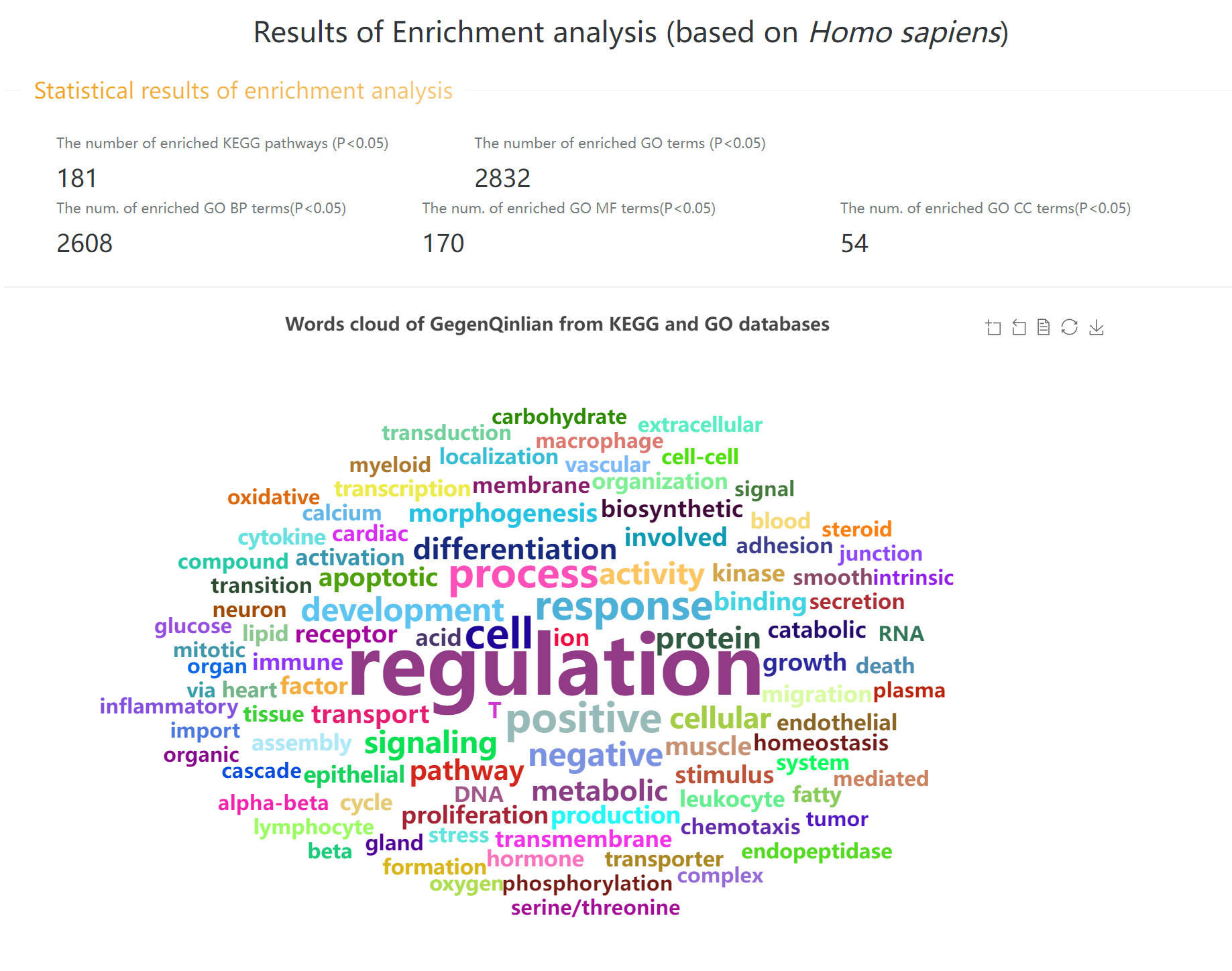

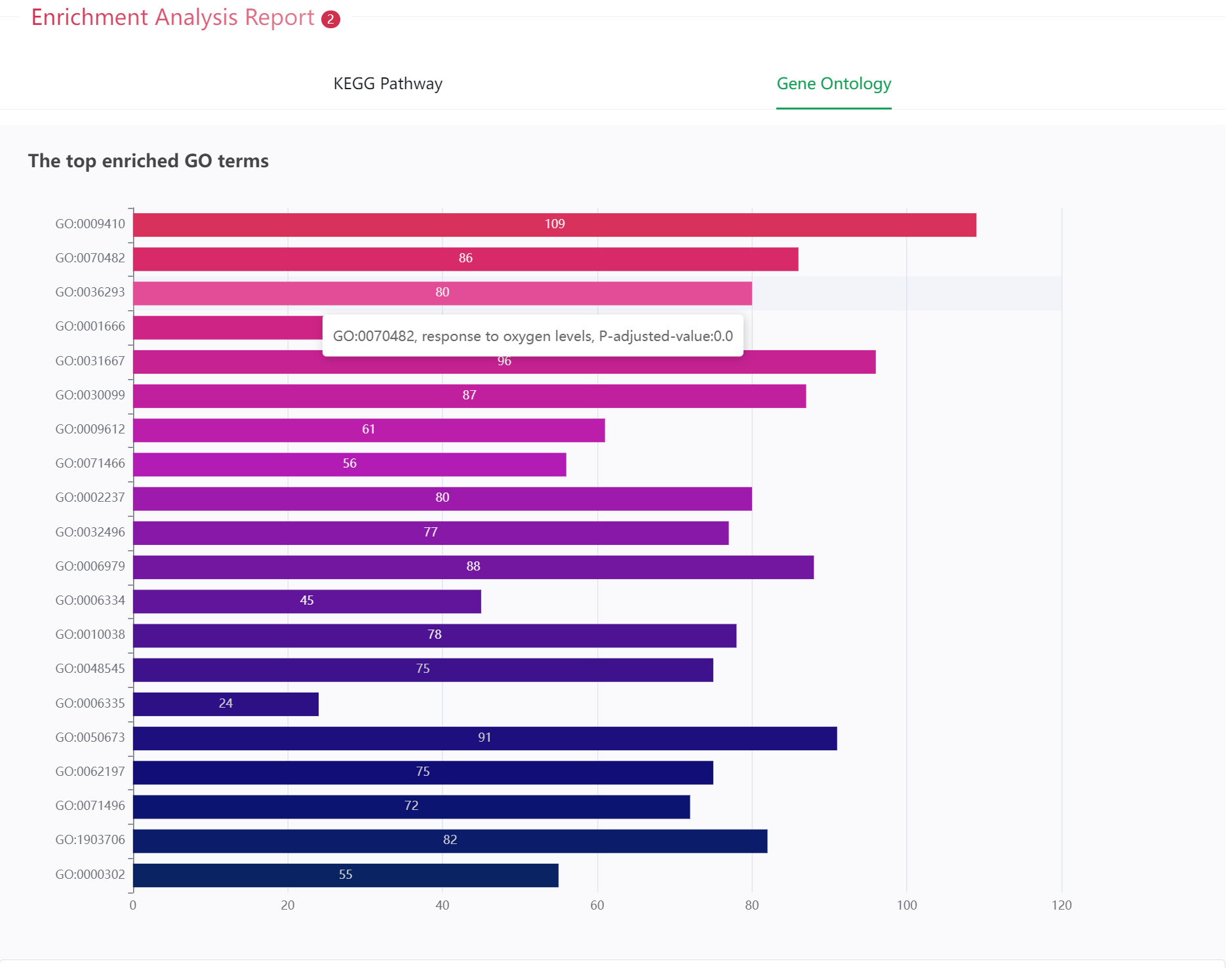

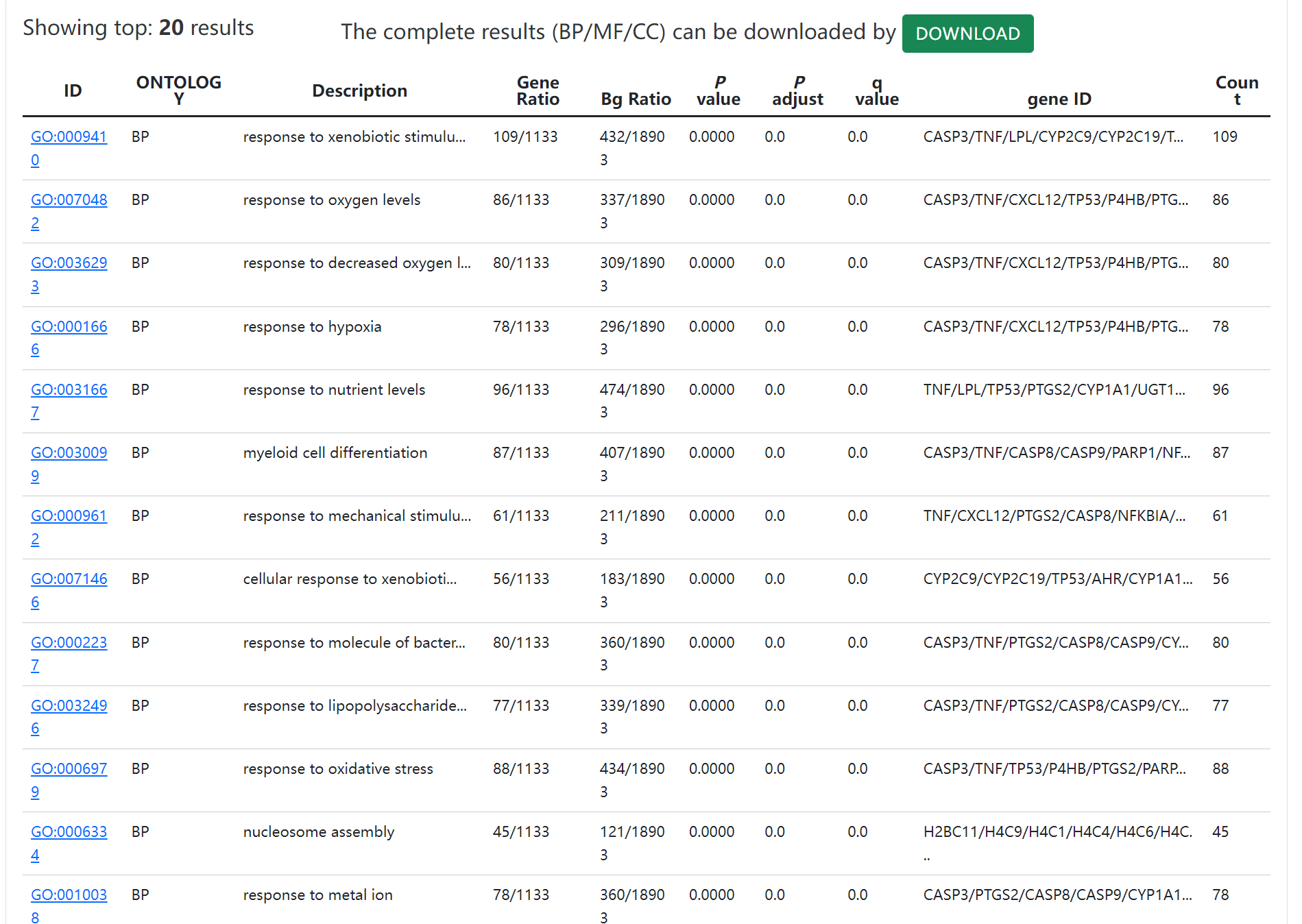
